# Nuclear damage in *LMNA* mutant iPSC-derived cardiomyocytes is associated with impaired lamin localization to the nuclear envelope

**DOI:** 10.1101/2021.10.30.466591

**Authors:** Melanie Wallace, Hind Zahr, Shriya Perati, Chloé D. Morsink, Lindsey E. Johnson, Anthony M. Gacita, Shuping Lai, Lori L. Wallrath, Ivor J. Benjamin, Elizabeth M. McNally, Tyler J. Kirby, Jan Lammerding

## Abstract

The *LMNA* gene encodes the nuclear envelope proteins Lamins A and C, which comprise a major part of the nuclear lamina, provide mechanical support to the nucleus, and participate in diverse intracellular signaling. *LMNA* mutations give rise to a collection of diseases called laminopathies, including dilated cardiomyopathy (*LMNA*-DCM) and muscular dystrophies. Although nuclear deformities are a hallmark of *LMNA*-DCM, the role of nuclear abnormalities in the pathogenesis of *LMNA*-DCM remains incompletely understood. Using induced pluripotent stem cell-derived cardiomyocytes (iPSC-CMs) from *LMNA* mutant patients and healthy controls, we show that *LMNA* mutant iPSC-CM nuclei have altered shape or increased size compared to healthy control iPSC-CM nuclei. The *LMNA* mutation exhibiting the most severe nuclear deformities, R249Q, additionally caused reduced nuclear stiffness and increased nuclear fragility. Importantly, for all cell lines, the degree of nuclear abnormalities corresponded to the degree of Lamin A/C and Lamin B1 mislocalization from the nuclear envelope. The mislocalization was likely due to altered assembly of Lamin A/C. Collectively, these results point to the importance of correct lamin assembly at the nuclear envelope in providing mechanical stability to the nucleus and suggest that defects in nuclear lamina organization may contribute to the nuclear and cellular dysfunction in *LMNA*-DCM.

## Introduction

Lamin A/C are intermediate filaments that assemble to form the nuclear lamina, a dense protein meshwork that resides underneath the inner nuclear membrane. The nuclear lamina provides structural support to the nucleus and functions in diverse mechanical and biochemical signaling processes (Maurer and Lammerding, 2019; Donnaloja *et al*., 2020; Wang *et al*., 2022a; Shah *et al*., 2023). More than 450 mutations have been identified in the *LMNA* gene that give rise to a collection of about 15 different human diseases, termed ‘laminopathies,’ which include dilated cardiomyopathy (*LMNA*-DCM), Emery-Dreifuss muscular dystrophy (EDMD), congenital muscular dystrophy (CMD), and Hutchinson-Gilford progeria syndrome (HGPS) (Davidson and Lammerding, 2014). *LMNA*-DCM has a high mortality rate, and compared to other forms of congenital DCM, *LMNA-*DCM has a particularly poor prognosis with early onset, a high occurrence of arrhythmias, and up to 19% of all patients requiring heart transplants (Taylor *et al*., 2003; McNally and Mestroni, 2017; Hasselberg *et al*., 2018). To date, clinical treatments for *LMNA*-DCM have focused on slowing heart failure rather than targeting the underlying cellular pathology, largely because the molecular disease mechanisms responsible for *LMNA*-DCM are not yet fully understood.

*LMNA* mutations and deletions often cause a reduction in nuclear stiffness that results in changes to nuclear shape, nuclear envelope rupture, and altered response to mechanical stress (Broers *et al*., 2004; Lammerding *et al*., 2004; Nikolova *et al*., 2004; Chandar *et al*., 2010; De Vos *et al*., 2011; Zwerger *et al*., 2013; Cho *et al*., 2019; Bertrand *et al*., 2020; Earle *et al*., 2020). However, few studies have been completed to understand nuclear mechanics and nuclear deformation in *LMNA* mutant human cardiomyocytes. Until the recent advent of human induced pluripotent stem cell (iPSC) models, the majority of studies on the cellular disease pathogenesis of *LMNA*-DCM have relied on mouse models that do not fully reflect all aspects of the human disease. For example, mutant mouse phenotypes require homozygous mutant alleles rather than dominant heterozygous allelic state in human disease (Stewart *et al*., 2007). As such, limited previous studies have demonstrated nuclear shape changes and nuclear envelope ruptures resulting in the mislocalization of cellular components into the nucleus in cardiomyocytes of laminopathy models (Nikolova *et al*., 2004; Chandar *et al*., 2010; Cho *et al*., 2019; Shah *et al*., 2019). Despite these efforts, the cardiac-specific mechanisms through which nuclei become damaged, the consequences of nuclear damage, and the effect of specific *LMNA* mutations in the context of nuclear damage in cardiomyocytes remain largely unclear. Since nuclear stability is necessary to resist the mechanical stress from contractile forces in cardiomyocytes (Swift *et al*., 2013; Buxboim *et al*., 2014; Cho *et al*., 2019; Heffler *et al*., 2019; Piccus and Brayson, 2020; Leong *et al*., 2023), it is critical to determine the cardiomyocyte-specific role of nuclear mechanics and damage in *LMNA*-DCM and how they contribute to disease progression.

Here, we use three *LMNA* mutant patient-derived human iPSC lines, iPSCs from healthy controls, and iPSC-derived cardiomyocytes (iPSC-CMs) to provide a quantitative analysis of the causes and consequences of nuclear damage in *LMNA*-DCM, with a particular focus on the nuclear lamina organization and physical properties of the cell nucleus. We show that different *LMNA* mutant iPSC-CMs exhibit varying degrees of nuclear shape and size deformities, with only cells with the most severe mutation, *LMNA* R249Q, exhibiting nuclear envelope rupture. Furthermore, we demonstrate that the severity of nuclear deformities correlates with reduced Lamin A/C and Lamin B1 localization to the NE, likely due to defective lamin assembly. Assembly of Lamin A/C at the nuclear envelope is critical for maintaining nuclear mechanical properties and nuclear envelope integrity (Zwerger *et al*., 2015; Cho *et al*., 2019; Earle *et al*., 2020). As such, defective assembly of lamins at the nuclear envelope may cause reduced nuclear stability and/or defects in organization of nuclear envelope proteins that in turn may result in altered cytoskeletal organization, abnormal nuclear shape, and susceptibility of the nucleus to damage. Collectively, these findings suggest that mislocalization of Lamin A/C and Lamin B1 from the nuclear envelope may explain nuclear deformities and damage observed in three *LMNA*-DCM iPSC-CM lines.

## Materials and Methods

### iPSC culture and cardiac differentiation

Healthy control (WT) and patient-derived *LMNA* mutant iPSCs were either purchased from commercial sources (Cure CMD) or generously provided from other researchers (Figure 1A; Table 1). iPSCs were cultured feeder-free in TeSR-E8 media (#05990, StemCell Technologies) on 1:30 Matrigel Basement Membrane (Matrigel; #47743-722, Corning) diluted in RPMI 1640 medium (#11875093, Gibco). Media was changed daily, and colonies were monitored for and removed with signs of spontaneous differentiation.

**Figure 1.**
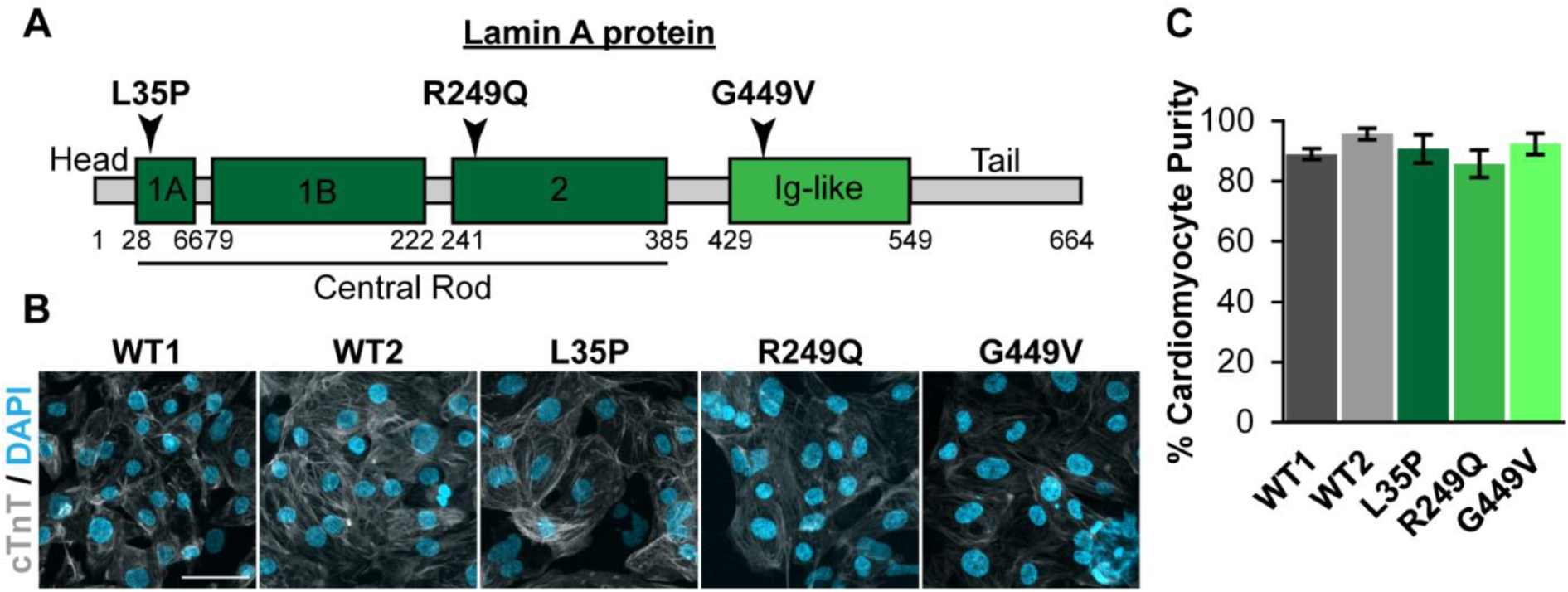
*LMNA* mutant iPSCs differentiate into cardiomyocytes with high efficiency. (A) Diagram of Lamin A protein domains with the locations of the amino acid substitutions used in this study indicated, which map to separate domains of the Lamin A protein. Note that these domains are also present in the Lamin C protein, which has an alternative tail domain compared to Lamin A. (B) Immunofluorescence images of iPSC-CMs show expression of the cardiac marker Cardiac Troponin T (cTnT) in the vast majority of cells. Scale bar = 50 μm. (C) Quantification of iPSC-CM purity based on cTnT labeling shows that all cell lines differentiate into cardiomyocytes with >85% efficiency, with no statistically significant differences between the cell lines. Data represented as mean ± SEM. *N* = 8-14 images per group.

**Table 1.** Overview of LMNA-mutant iPSC-CM lines. The iPSC lines were isolated from patients carrying LMNA mutations that produce amino acid substitutions in different domains of the Lamin A/C protein, as indicated in the table, and cause either Congenital Muscular Dystrophy (CMD) or LMNA-DCM. The CMD patients with the L35P and G449V amino acid substitutions are expected to develop a cardiac phenotype later in life, in addition to the skeletal muscular dystrophy present at diagnosis. *The G449V patient presented with muscular dystrophy in childhood, but patient is too young to necessarily develop cardiac phenotype.

| Protein | cDNA | Exon | Protein Domain | Mutation Origin | Affected Tissue | Cell Source |
| --- | --- | --- | --- | --- | --- | --- |
| **p.L35P** | c.T104C | 1 | Coil 1A | Unknown | Skeletal and cardiac muscle | Cure CMD and Cellular Dynamics International Inc. |
| **p.R249Q** | c.G746A | 1 | Coil 2 | De novo | Skeletal and cardiac muscle | Elizabeth McNally |
| **p.G449V** | c.G1346T | 7 | Ig-fold | De novo | Skeletal muscle* | Ivor Benjamin |

Cardiac differentiation was performed based on a previously established protocol (Sharma *et al*., 2015). iPSCs were washed with RMPI 1640 medium, dissociated into single cells with TrypLE Express (#12605028, Gibco), and seeded on 1:30 Matrigel in 12-well plates in TeSR-E8 media supplemented with Y-27632 (10 μM, #10005583, Cayman Chemical). Media was changed to fresh TeSR-E8 after 24 hours. After 48 hours (D0), differentiation was induced using inhibition of the WNT pathway with CHIR99021 (6 μM or 12 μM, #SML1046, Sigma-Aldrich) in RPMI 1640 media supplemented with B27 minus insulin (#A1895602, Gibco; referred to here as B27-INS). The CHIR99021 concentration, at 6 μM or 12 μM, was optimized for each cell line as each had different sensitivities to the inhibitor. Media was changed to fresh B27-INS at day 1 of differentiation (D1). IWR-1 (5 µM, #13659, Cayman Chemical) in B27-INS was added at D3 and changed to fresh B27-INS at D5. Media was changed to RPMI 1640 medium supplemented with B27 (#17504044, Gibco; referred to as B27) on D7 and was changed daily for three days until beating was induced. On D10, media was changed to RPMI 1640 without L-Glutamine (#21870076, Gibco; referred to as selection medium) supplemented with B27 and Sodium Lactate (Sigma) for metabolic selection of cardiomyocytes. On D12, media was changed to B27 supplemented with Penicillin-Streptomycin (referred to as B27+P/S), and iPSC-CMs were allowed to recover for two days before passaging (‘first CM passage’).

At D14, only wells of iPSC-CMs that were beating were washed with RPMI 1640 and dissociated into single cells using Trypsin (0.25%). Trypsin was inactivated using RPMI 1640 plus 20% fetal bovine serum. Cells were centrifuged and resuspended in RPMI 1640 plus fetal bovine serum (10%) and 10 μM Y-27632 and then seeded 1:1 in 12-well plates coated with Fibronectin (4 μM). After 24 hours, media was changed to B27+P/S. After an additional 24 hours, iPSC-CMs were subjected to a second round of metabolic selection with selection medium for 48 hours. Media was then changed back to B27+P/S, and cells were used seven days after the first CM passage (D21).

After 21 days of differentiation, all cell lines spontaneously contracted throughout the wells. We confirmed cardiac differentiation by immunofluorescence staining for a cardiac marker, cardiac troponin T (cTnT; Figure 1B). We quantified the percentage of cells that expressed cTnT, which demonstrated that all cell lines routinely had over 85% expression of cTnT and no significant changes in expression between cell lines (Figure 1C). All immunofluorescence experiments were performed with a cTnT co-stain. Upon immunofluorescence labeling, only wells with predominantly cTnT-expressing cells were used for analysis (wells with few cTnT-expressing cells were excluded), and any remaining individual cells not expressing cTnT were excluded from analysis.

### Generation of fluorescently labeled cell lines

iPSC-CMs after the first CM passage were stably modified with lentiviral constructs to express a nuclear envelope rupture reporter consisting of a green fluorescent protein with a nuclear localization sequence (NLS-GFP, full vector: pCDH-CMV-NLS-copGFP-EF1-blastiS) described previously (Denais *et al*., 2016). Cells were cultured for at least six days after modification to allow sufficient expression of the construct. Expression of NLS-GFP was confirmed before each experiment.

### Lamin A/C depletion

Prior to the first round of metabolic selection, WT2-iPSC-CMs were stably modified with a lentivirus to express either shRNA targeting *LMNA* (shLMNA; pLKO.1-shRNA-mLMNA; target sequence: CCGGGAAGCAACTTCAGGATGAGATCTCGAGATCTCATCCTGAAGTTGCTTCTTTTTG) or a non-target control (shNT). For all experiments, shRNA-modified iPSC-CMs were used as a mixed population of modified (‘knockdown’ or ‘KD’) and un-modified (‘no knockdown’ or ‘no KD’) cells to serve as an additional internal control group, as described in the Image Analysis section.

### Long-term imaging experiments

Long-term fluorescence imaging of iPSC-CMs expressing NLS-GFP imaging was performed on an IncuCyte (Sartorius) incubator imaging system. Images were acquired at 10× magnification every 15 minutes for three days, starting one week after the first CM passage. Images were exported and manually analyzed in FIJI software to quantify the frequency and duration of nuclear envelope rupture, as evidenced by the transient loss of NLS-GFP from the nucleus.

### Immunofluorescence staining

iPSC-CMs were passaged one week after the first CM passage into optically clear 96-well plates. The next day, media was changed to B27+P/S for several hours before cells were washed with 1× PBS and fixed in warm Paraformaldehyde (4%) for 10 minutes. Cells were then washed with 1× PBS and blocked in 3% BSA with 0.1% Triton-X 100 (Thermo-Fisher) and 0.1% Tween (Sigma) in PBS for one hour at room temperature. Primary antibodies (Table S1) were prepared in blocking solution and incubated overnight at 4°C. The Lamin A/C antibody used (Santa Cruz; E-1) targeted the N-terminus (AA 2-29) to stably detect both the Lamin A and C isoforms. The Lamin B antibody used (proteintech; 12987-1-AP) targets only Lamin B1. Following primary antibody incubation, iPSC-CMs were then washed with a solution of 0.3% BSA with 0.1% Triton-X 100 and 0.1% Tween in PBS and stained with AlexaFluor secondary antibodies (1:250, Invitrogen) for 1 hour at room temperature. DAPI (1:1000, Sigma) was added for 15 minutes at room temperature, and cells were washed with PBS before imaging.

### Contractility measurements

iPSC-CMs were plated at a concentration of 400,000 cells per well on Matrigel Basement Membrane (Matrigel; #47743-722, Corning)-coated 24-well black 14 mm plates (IBIDI, 82421) in RPMI supplemented with B27+P/S (11875093, 17504044 and 15140122, Thermofisher). Media was changed every other day. Cells contractility was measured around day 7 after seeding, when the iPSC-CMs exhibited robust contractions. Contractility measurements were performed without pacing using the Cytocypher Multicell High Throughput System (Cytocypher BV, IonOptix Corporation, Westwood, MA, United States). Measurements were performed in a temperature-controlled chamber with a temperature of 36.5°C and CO_2_ levels of 5%. Cells that were spontaneously beating were used for measurements. Each cell was measured for 10 seconds, measuring 10 contraction traces. For each replicate, measurements of 12-15 contracting cells were collected. Data was analyzed with the Transient Analysis Tool software (Cytocypher BV). The following calculations were performed to obtain contraction and relaxation times:

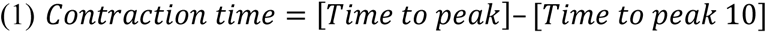

and

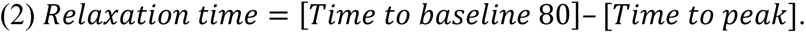

In these equations, *Time to peak* is the time it takes iPSC-CM to reach maximum contraction; *Time to peak 10* is the time to reach the initial 10% of maximum contraction; *Time to baseline* is the time it takes iPSC-CM to relax after contractions and return to their original length; *Time to baseline 80* is the time it takes to reach 80% of full relaxation.

### Pharmaceutical treatments

To determine the role of actin and microtubules in R249Q mutant iPSC-CM nuclear deformities, WT2 and R249Q iPSC-CMs were passaged one week after the first CM passage into optically clear 96-well plates. The next day, iPSC-CMs were treated for one hour with drugs to stabilize or depolymerize microtubules (10 nM Paclitaxel or 0.5 µg/mL Nocodazole, respectively) or to depolymerize or inhibit contractility of actin (1 μM Latrunculin B or 25 μM Blebbistatin, respectively), or DMSO as a vehicle control. Cells were then washed with 1× PBS and immunofluorescently labeled as specified in the text.

### Nuclear semi-permeabilization and soluble protein washout

One week after the first CM passage, iPSC-CMs were passaged into optically clear 96-well plates (Greiner). The next day, media was changed to B27+P/S for several hours before cells were washed with 1× PBS. A buffer, “CSKT,” containing NaCl (100 mM), Sucrose (300 mM), MgCl_2_ (3 mM), PIPES (10 mM, pH 6.8) and Triton-X 100 (0.5%) was added to cells to semi-permeabilize nuclei and incubated on ice for 1 minute. A 1-minute wash with 1× PBS was used as a control in parallel to semi-permeabilization with CSKT. CSKT buffer was removed, cells were gently washed with 1× PBS to washout soluble nuclear proteins, and then the cells were fixed with 4% PFA for 10 minutes. Following PFA fixation, cells were inspected under a microscope to ensure that nuclear semi-permeabilization and washout was successful, as evidenced by the darkened and more prominent appearance of nuclei.

### Image acquisition

Plates were imaged on an inverted Zeiss LSM700 confocal microscope. Z-stacks were taken using a 40× water immersion (1.2 NA) objective with Airy units set to 1.0 for all images. Z-stacks for nuclear volume were acquired using the optimum step size for the 40× water immersion objective, i.e., 0.35 µm. All other image stacks were acquired using a step size of 1.5 µm and analyzed using Maximum Intensity Projections (MIPs).

### Image analysis

All image analysis was conducted by observers blinded for genotype and treatment conditions. iPSC-CM purity was quantified from MIPs by counting the proportion of nuclei in the image that are located within cardiac troponin T (cTnT) positive cells (referred to as cTnT-positive nuclei). For image analysis of iPSC-CMs, only cTnT-positive nuclei were used.

For nuclear shape and area, a custom MATLAB (MathWorks) script was used to segment nuclei based on the Lamin B1 immunofluorescence signal in MIPs. Nuclei intersecting with the image edges were removed, followed by manual selection of nuclei in each image to exclude any nuclei touching each other, dead cells, or non-cTnT-positive nuclei. The area (A) and perimeter (P) were then measured for each selected nucleus, and the circularity index (CI) was computed based off the formula 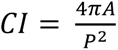. A perfectly circular object has a circularity index of 1, and the circularity index decreases as circularity decrease, e.g., in oval or irregularly shaped nuclei.

Nuclear volume was computed from high resolution confocal Lamin B1 immunofluorescence 3D image stacks. Image stacks were read into the FIJI image analysis program, and a threshold value was determined from the middle slice of z-stacks. The threshold value was then input into the 3D Simple Segmentation suite (Ollion *et al*., 2013), and the 3D Objects Counter (Bolte and Cordelières, 2006) was used to compute nuclear volumes. Partial nuclei at the image edges, nuclei touching each other, dead cells, and non-cTnT-positive nuclei were excluded from analysis.

For shRNA experiments, nuclear area, circularity index, and Lamin A/C mean fluorescence intensity were computed from a modified version of the MATLAB script for nuclear area and circularity described above. Nuclear area and circularity index were computed from the Lamin B1 immunofluorescence signal of MIPs as described above. Additionally, Lamin A/C images were read into the program, images were cross-correlated with the corresponding Lamin B1 images to analyze only the user-selected nuclei, and the mean Lamin A/C immunofluorescence intensity inside of the area of each user-selected nucleus was computed. For each individual experiment, Lamin A/C intensities for shNT- and shLMNA-treated cells were plotted, and the lowest Lamin A/C intensity from the shNT group was used as a threshold, above which shLMNA nuclei were considered “no knockdown (no KD)” and below which shLMNA nuclei were considered “knock down (KD).” shLMNA nuclei were separated into “no KD” or “KD” groups for each experiment, with “no KD” nuclei serving as a secondary, internal control. Nuclear area and circularity index were analyzed based on the average of several separately thresholded experiments.

Nuclear volume analysis of shRNA Lamin A/C depleted cells and controls was performed using the same FIJI image analysis pipeline described above. For these experiments, an observer blinded for treatment conditions manually classified nuclei as “no KD” or “KD” for each image based on their Lamin A/C immunofluorescence intensity. Untreated control or shNT groups, as expected, had extremely few nuclei considered as “KD” and the proportion of shLMNA no KD and KD nuclei was similar to that obtained by the objective thresholding method.

Fluorescence intensity profiles were computed from high-resolution confocal z-stacks in the ZEN software (Zeiss). For each nucleus, the z-position for which the x-y plane dissects the center of the nucleus was identified, and a line was drawn across the major axis of the nucleus to obtain the Lamin A/C and Lamin B1 fluorescence intensity values along with distances. Fluorescence intensity profiles were trimmed such that the first values in the profile correspond to the outer edges of the NE. Distances across each nucleus were then normalized to a scale of 0 to 1 to account for differences in nuclear size. Fluorescence intensity profiles and normalized distances for each nucleus for each cell line were then read into a custom R script for analysis (script available upon request). Fluorescence intensity profiles across all nuclei in a cell line were averaged and normalized to the area under the curve to account for variations in fluorescence intensities between cell lines due to varying lamin levels or staining conditions. The boundary of the nuclear lamina was determined by the inflection point of the average fluorescence intensity profile of all cell lines with the nuclear lamina defined as the peripheral regions with normalized nuclear distance of <0.16 and >0.84. For each nucleus, the fluorescence intensity of nucleoplasmic lamins was then determined by taking the 95^th^ percentile of fluorescence intensity values in the nuclear lamina region (to avoid influence of spurious pixel intensity values), and the fluorescence intensity of nucleoplasmic lamins was determined by taking the average value of fluorescence intensities in the nucleoplasm, corresponding to normalized nuclear distances between 0.16 and 0.84. The ratio of nucleoplasmic to peripheral lamins was then computed for each nucleus.

Lamin A/C and phospho-Lamin A/C fluorescence intensities were computed using the same MATLAB script used for shRNA analysis described above with Lamin A/C images being used for thresholding and user selection under the same exclusion criteria as previously selected. All Lamin A/C and phospho-Lamin A/C average fluorescence intensity values were normalized to the respective average of a healthy control (WT1) fluorescence intensity from images taken at the same time.

### Nuclear stiffness measurements

iPSC-CMs were passaged one week after the first CM passage into glass-bottom 35-mm tissue culture plates (FluoroDish; World Precision Instruments) coated with 4 µM Fibronectin and allowed to adhere for 24 hours. Prior to experimentation, nuclei were fluorescently labeled with Hoechst 33342 (1:1000; Biotium). Experiments were performed at room temperature, and all indentations were performed within one hour of removing cells from the incubator, at which point the majority of cells were still beating.

Nanoindentation was performed with a microscope-mounted Chiaro system (Optics11). The system was calibrated prior to experimentation according to the manufacturer’s instructions, and a spherical glass tip with a radius of 3 µm and stiffness of 0.026 N/m was used to indent cells. The center of the probe tip and the center of nuclei were aligned, and an image of each fluorescent nucleus was taken prior to indentation. The probe was then lowered to approximately 5-10 µm above the top of the nucleus. If the probe accidentally touched a nucleus prior to the start of the indentation, the cell was excluded. The probe was lowered towards the nucleus at a speed of 2 µm/s, and once the system reached a contact force threshold of 5 nN, indentation continued at a rate of 10 nN/s until the probe reached a force threshold of 40 nN. The 40 nN load was held for five seconds, and then the probe was retracted at a rate of 10 nN/s. A Hertzian model with a Poisson’s Ratio of 0.5 (Guz *et al*., 2014) was fitted to the load vs. indentation curve from 250-1500nm of indentation depth for each nucleus to determine the Young’s elastic modulus. Any nucleus with a fit for the Hertzian model below an R^2^ value of 0.95 was excluded from further analysis.

### Western analysis

iPSC-CMs lysates were collected one week after the initial passage. Cells were lysed in High Salt RIPA buffer with protease (cOmplete EDTA-Free, Roche), phosphatase inhibitors (PhosSTOP, Roche) and protease inhibitor Phenylmethanesulfonyl fluoride (PMSF, Sigma-Aldrich). After lysis, samples were vortexed for five minutes, sonicated for 30 seconds at 36% amplitude, boiled for five minutes at 93°C, and sheared with a syringe. Protein content was quantified with Bio-Rad Protein Assay Dye, and 20 µg of protein was run on a 4-12% Bis-Tris polyacrylamide gel using an SDS-PAGE protocol. Protein was transferred to a polyvinylidene fluoride (PVDF) membrane using a semi-dry transfer at 16 V for one hour. Membranes were blocked with Intercept (PBS) Blocking Buffer (LI-COR) for one hour at room temperature, primary antibodies were diluted in the same blocking buffer, and membranes were incubated the primary antibodies on a rocker overnight at 4°C. IRDye 680LT or IRDye 800CW (LI-COR) secondary antibodies were used to detect protein bands. Membranes were imaged on an Odyssey® CLx imaging system (LI-COR) and analyzed using Image Studio Lite (LI-COR).

### Gene expression analysis

Total RNA was isolated one week after the initial passage using Trizol/Chloroform extraction and RNeasy Mini kit (Qiagen) purification. cDNA was synthesized from 1 μg of total RNA using the iScript cDNA synthesis kit (Bio-Rad). qPCR was performed using the LightCycler 480 (Roche) and LightCycler 480 SYBR Green I Master Mix (Roche) using the following parameters: an initial denaturation step of 95°C for 10 min, 50 cycles of (95°C for 15s, 60°C for 30s, 72°C for 30s), and a final cooling cycle of 4°C for 30s. Relative gene expression of *LMNA* and *LMNB1* genes was calculated using the Ct method and was normalized to the geomean of *GAPDH*, *18S* and *TMX4* housekeeping genes. The primer sequences used are provided in Supplementary Table 2.

### Code availability

Custom MATLAB scripts used in the analysis are available upon request.

### Statistical analysis

All results were taken from a minimum of three independent experiments. For continuous numeric datasets in which individual nuclei were analyzed, a mixed-effects linear regression was performed in R to account for the effects of both *LMNA*-mutations and individual cell lines. Datasets were tested for normality, and data not following a normal distribution were linearized either by taking the natural log or the square root, whichever achieved better normalization. Pairwise *t*-tests were then performed using the Tukey method to correct for multiple comparisons. For continuous numeric datasets in which whole images or experiments were analyzed and followed a normal distribution, either a Student’s *t*-test (for two groups) or one-way analysis of variance (ANOVA) with multiple comparisons (for more than two groups) was performed in Prism (GraphPad). Dunnett correction was used for multiple comparisons. For nuclear envelope rupture experiments in which proportional data was analyzed, a binomial regression was performed in R. Since several cell lines exhibited no nuclear envelope rupture, a Firth logistic regression was then performed to reduce bias from such zero values. Pairwise *t*-tests were then performed using the Tukey method to correct for multiple comparisons. For correlation data, a linear regression was performed for each cell line in GraphPad Prism, and the correlation was tested for significance for a non-zero slope using Pearson correlation and compared to the other regression. Unless otherwise noted, error bars in graphs represent mean ± standard error of the mean (SEM).

## Results

### LMNA mutant iPSCs differentiate efficiently into cardiomyocytes

To quantitatively study the effects of different *LMNA* mutations on nuclear lamina organization and nuclear mechanics, we selected three *LMNA* mutant iPSC lines (*LMNA* L35P, R249Q, and G449V) derived from patients with DCM and CMD and two healthy controls (WT1, WT2) for this study (Table 1). Each of the *LMNA* mutations cause amino acid substitutions in a different domain of the Lamin A/C protein (Figure 1A), enabling the identification of common and mutation-specific effects. The iPSCs were subjected to a chemically defined cardiac differentiation protocol to obtain iPSC-derived cardiomyocytes (iPSC-CMs). All iPSCs underwent successful cardiac differentiation with over 85% of nuclei staining positive for cardiac troponin T (cTnT), an established cardiomyocyte marker (Figure 1B, C). We did not detect any significant differences in iPSC-CM differentiation efficiency between cell lines (Figure 1C).

### LMNA mutant iPSC-CMs express reduced levels of Lamin A/C protein

Although most *LMNA* mutations result in stable expression of the mutant protein, some *LMNA* mutations can reduce protein stability and lead to haploinsufficiency (Wolf *et al*., 2008; Siu *et al*., 2012; Cattin *et al*., 2013). To test whether the *LMNA* mutant cell lines had altered expression of Lamin A/C, we investigated protein and mRNA expression levels of Lamin A/C and Lamin B1 in the panel of iPSCs and the corresponding iPSC-CMs. We detected only very low Lamin A/C protein levels in iPSCs by both western analysis (Figure 2A) and immunofluorescence labeling (Figure S1A), consistent with previous reports that iPSCs express no or only little Lamin A/C (Zuo *et al*., 2012). As expected, *LMNA* mRNA and Lamin A/C protein expression increased upon differentiation into iPSC-CMs (Figure 2A, Figure S2A). Although we did not detect differences in *LMNA* mRNA expression among the iPSC-CM cell lines (Figure S2A), Lamin A/C protein expression varied substantially between healthy control and *LMNA*-mutant iPSC-CMs. Both the L35P and R249Q mutant iPSC-CMs had significantly decreased Lamin A/C protein expression compared to either one or both healthy control cell lines (Figure 2A-B, Figure S2C-D, Figure S3A), suggesting differences in translation or protein turnover of the mutant Lamin A/C. These data further indicate that reduced protein level of Lamin A/C in L35P and R249Q mutant iPSC-CMs may result in partial haploinsufficiency. By contrast, Lamin B1 was present at high levels in iPSC and iPSC-CMs, with the L35P iPSC-CMs showing increased protein levels compared to the other iPSC-CM lines, despite similar *LMNB1* mRNA expression (Figure 2A-B, Figure S2B).

**Figure 2.**
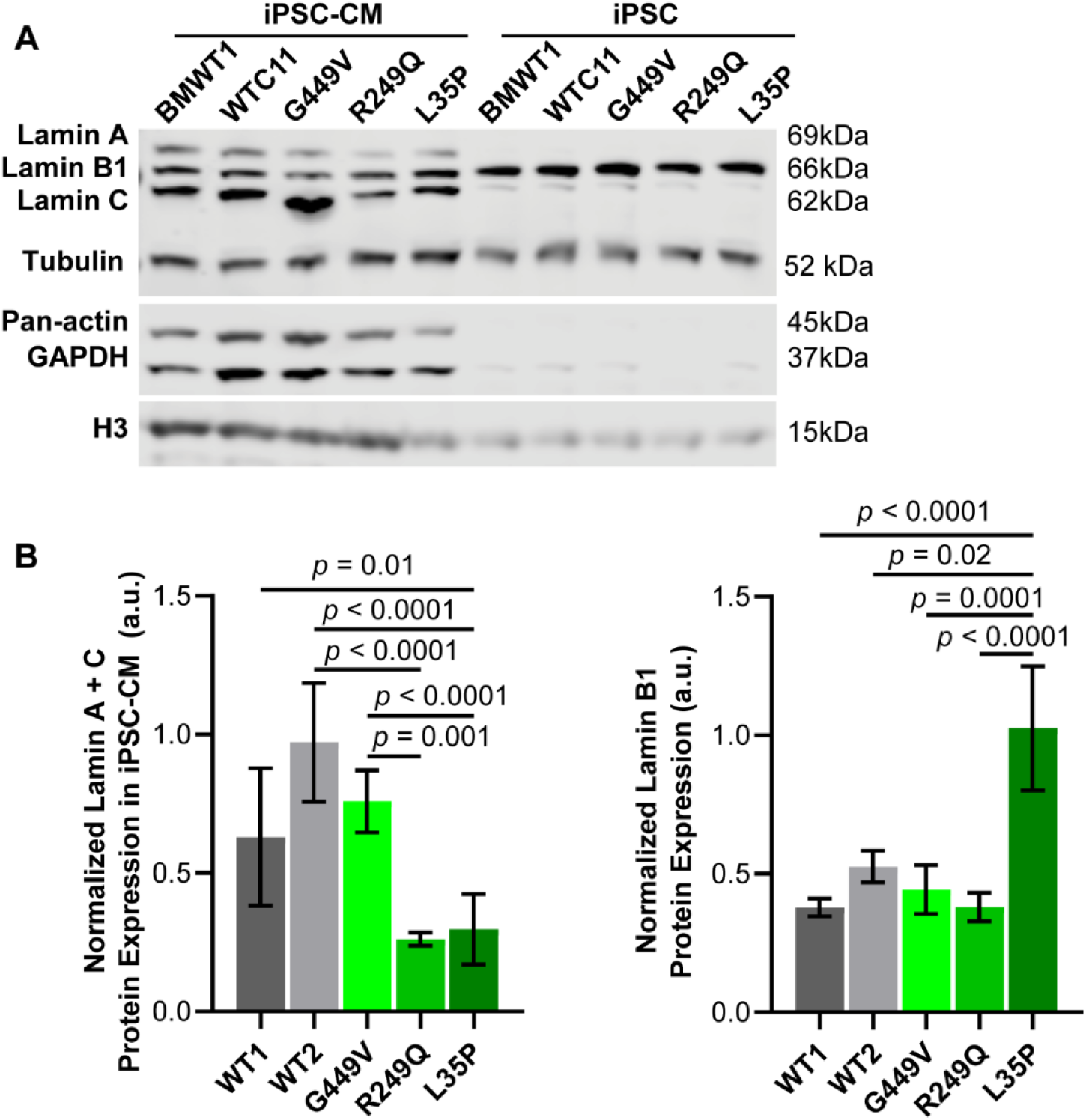
R249Q and L35P iPSC-CMs have reduced Lamin A/C protein expression. (A) Representative western analysis, showing that iPSCs express only barely detectable Lamin A/C, and that Lamin A/C expression increases in iPSC-CMs. (B) Quantification of Lamin A/C protein expression from western analysis of iPSC-CM samples, showing that both WT and *LMNA-*mutant iPSC-CMs have variations in levels of Lamin A/C. Only L35P iPSC-CMs had a significant decrease in Lamin A/C protein compared to both healthy controls, whereas R249Q iPSC-CMs had significantly decreased Lamin A/C protein compared to only WT2 iPSC-CMs. (C) Quantification of Lamin B1 protein expression shows that L35P iPSC-CMs had significantly increased expression of Lamin B1, whereas all other cell lines had similar expression levels. Data presented as mean ± SEM. *N* = 6 protein lysates per group.

### LMNA mutant iPSC-CMs exhibit altered nuclear shape and size

Although altered nuclear shape and size are hallmarks of skeletal muscle and cardiac laminopathies, the severity of nuclear shape defects varies substantially across different *LMNA* mutations (Sullivan *et al*., 1999; Lammerding *et al*., 2004, 2006; Muchir *et al*., 2004; Zwerger *et al*., 2013; Steele-Stallard *et al*., 2018; Bertrand *et al*., 2020). To quantify defects in nuclear morphology in the different *LMNA* mutant iPSC-CMs, we computed nuclear circularity index, area, and volume based on confocal 3D image stacks and cross-sections of iPSC-CMs immunofluorescently labeled for Lamin B1 (Figure 3A). The circularity index has a maximum value of one for perfectly circular nuclei and decreases in value for abnormally shaped nuclei (Figure 3B). R249Q mutant iPSC-CMs had a significantly decreased circularity index compared to healthy controls and other *LMNA* mutants (Figure 3C), indicating more abnormal nuclear shapes. In contrast, G449V mutant iPSC-CM nuclei exhibited no change in circularity index compared to healthy controls, and L35P mutant nuclei were slightly more circular compared to healthy controls and the other mutant iPSC-CMs (Figure 3C). Notably, all three *LMNA* mutant iPSC-CM lines had increased nuclear cross-sectional areas compared to healthy controls (Figure 3D), although only R249Q and L35P mutant iPSC-CMs showed reduction in nuclear height (Figure 3E) and increased nuclear volumes (Figure 3F) compared to healthy controls. These data suggest more severe nuclear defects in L35P and R249Q mutant iPSC-CMs with increased nuclear area and volume. These results may in part be explained by the reduced Lamin A/C protein expression in these cell lines, although we did not observe such trends in the WT1 iPSC-CMs, which had lower Lamin A/C levels compared to WT2 iPSC-CMs. The altered nuclear morphology in the *LMNA* mutant iPSC-CM lines points to potential changes in nuclear organization and/or nuclear stiffness, as decreased nuclear height and increased nuclear cross-sectional area can result in increased nuclear flattening under cytoskeletal tension.

**Figure 3.**
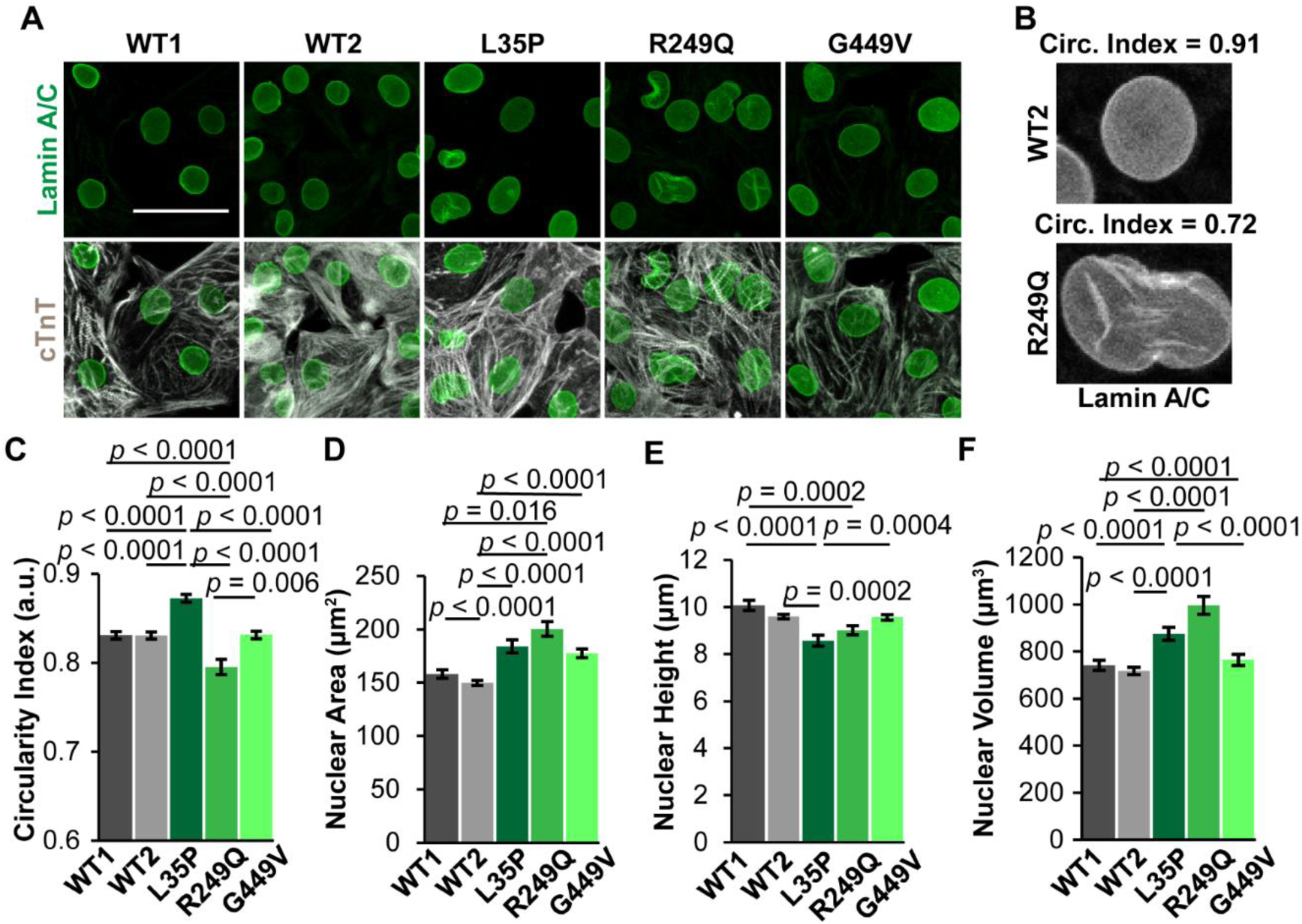
*LMNA* mutant iPSC-CMs have nuclear shape and size defects. (A) Representative immunofluorescence images of iPSC-CMs stained for Lamin A/C (green) and cardiac troponin T (cTnT, gray). Scale bar = 50 µm. (B) Examples of WT2 and R249Q nuclei with corresponding circularity index. (C) R249Q nuclei had significantly decreased circularity index values compared to the WT cell lines, whereas L35P and G449V nuclei had either an increase or no change in circularity index, respectively. (D) All *LMNA*-mutant iPSC-CMs exhibited an increase in nuclear area, but (E) only L35P and R249Q cells had decreased nuclear height. (F) L35P and R249Q iPSC-CMs had increased nuclear volumes. Data presented as mean ± SEM. *N* ≥ 83 nuclei per group for all experiments.

### R249Q-iPSC-CMs exhibit increased nuclear envelope rupture

In addition to changes to nuclear morphology, *LMNA* mutations can lead to increased incidence of nuclear envelope rupture due to weakening of the nuclear envelope (De Vos *et al*., 2011; Cho *et al*., 2019; Earle *et al*., 2020). Therefore, we introduced a nuclear envelope rupture reporter, NLS-GFP, in which a green fluorescent protein (GFP) is fused to a nuclear localization sequence (NLS)(Denais *et al*., 2016), into the iPSC-CMs. NLS-GFP normally localizes to the nucleus but leaks into the cytoplasm upon nuclear envelope rupture and then gradually translocates back into the nucleus following nuclear envelope repair (Figure 4A-B). Long-term (72 hours) time-lapse microscopy revealed that nuclear envelope rupture is extremely rare in healthy control iPSC-CMs (Figure 4C). In contrast, R249Q mutant iPSC-CMs, which had the most severe defects in nuclear shape, size, and volume (Figure 3C-E), had a significant increase of nuclear envelope ruptures compared to both healthy control cell lines (Figure 4C). On the other hand, the L35P and G449V mutant iPSC-CMs, which had milder defects in nuclear shape and size, showed no increase in nuclear envelope rupture or only a trend towards increased nuclear envelope rupture (G449V) that did not reach statistical significance (*p* > 0.3 versus healthy controls). Of note, the duration of nuclear envelope rupture was not statistically significant between any of the cell lines that exhibited nuclear envelope rupture (Figure S4), suggesting that *LMNA* mutations did not alter nuclear envelope repair, consistent with previous studies that found that Lamin A/C depletion did not alter nuclear envelope rupture duration (Denais *et al*., 2016; Halfmann *et al*., 2019). The dramatic increase of nuclear envelope rupture in the R249Q mutant iPSC-CMs compared to other *LMNA* mutant cell lines, paired with the R249Q iPSC-CMs being the only cell line to have a reduction in circularity index, suggests that R249Q mutant iPSC-CMs have more substantial changes in nuclear stability despite similar protein levels of Lamin A/C compared to WT1 and L35P iPSC-CMs.

**Figure 4.**
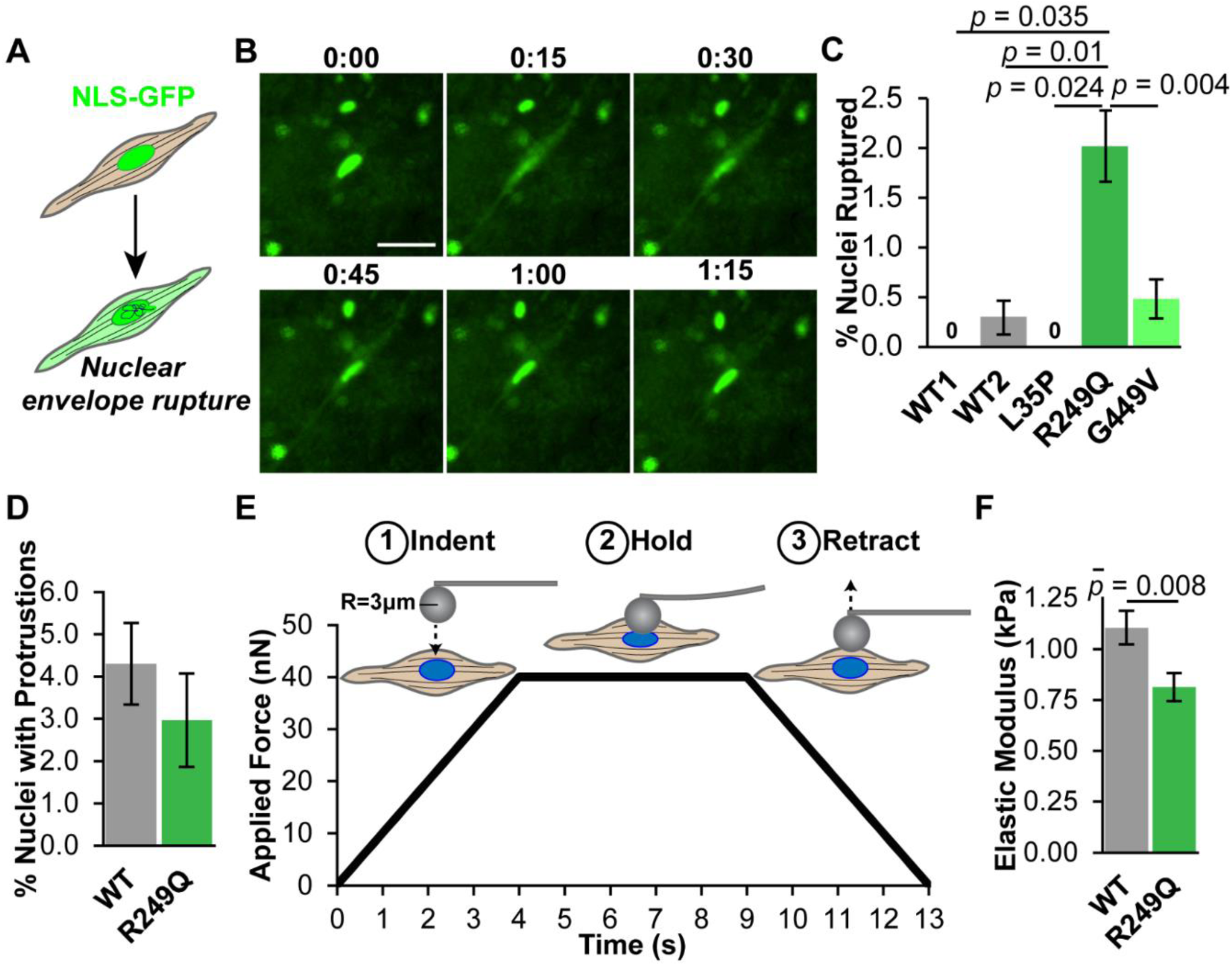
R249Q iPSC-CMs exhibit decreased nuclear stiffness and increased nuclear envelope rupture. (A) iPSC-CMs were modified with a nuclear envelope rupture reporter, NLS-GFP, consisting of GFP with a Nuclear Localization Sequence. NLS-GFP is normally packaged in the nucleus and leaks into the cytoplasm upon nuclear envelope rupture. (B) Representative time lapse series of a G449V iPSC-CM undergoing nuclear envelope rupture and repair. Scale bar = 30 µm. Time units are represented in hours:minutes. (C) Quantification of nuclear envelope ruptures shows that R249Q iPSC-CMs had significantly increased nuclear envelope rupture compared to the healthy control (WT) iPSC-CM lines. (D) R249Q iPSC-CMs have similar percentages of nuclei with chromatin protrusions to healthy control nuclei. (E) iPSC-CM nuclei were indented using a spherical probe on an Optics11 Chiaro nanoindenter to quantify nuclear stiffness. The indentation probe was lowered until it reached a load of 40 nN; the load was held for 5 seconds and then the probe was slowly retracted. (F) Quantification of nuclear elastic modulus, determined by fitting a Hertzian model to the load-indentation curves, revealed that R249Q nuclei are significantly softer compared to healthy control (WT) nuclei. Nuclear envelope rupture data and chromatin protrusions presented as mean ± SEP, nuclear stiffness data represented as mean ± SEM. *N* ≥ 1020 nuclei per group for nuclear envelope rupture, *N* ≥ 236 nuclei per group for chromatin protrusions, *N* ≥ 68 nuclei per group for nuclear stiffness.

Increased nuclear envelope rupture is often associated with formation of chromatin protrusions that extend beyond the nuclear lamina into the cytoplasm (Earle *et al*., 2020). Therefore, we quantified the frequency of chromatin protrusions in healthy control and R249Q iPSC-CMs. However, we did not observe any significant differences in the frequency of chromatin protrusions between healthy control and R249Q iPSC-CMs (Figure 4D).

### Actin and microtubules play a minor role in nuclear damage in R249Q iPSC-CMs

Nuclear abnormalities and nuclear envelope rupture arise from cytoskeletal forces on the nucleus (Hatch and Hetzer, 2016; Xia *et al*., 2018; Cho *et al*., 2019; Heffler *et al*., 2019; Earle *et al*., 2020). Furthermore, some *LMNA* mutations have been reported previously to affect iPSC-CM contractility (Lee et al., 2017; Miura et al., 2022; Perea-Gil et al., 2022). Therefore, to investigate if the increased nuclear defects in *LMNA* mutant iPSC-CMs result from altered cytoskeletal forces on the nuclei, we measured the contractility of the three *LMNA* mutant and two healthy control iPSC-CM lines. Quantification of spontaneous contractility of individual iPSC-CMs (Figure S5A), however, did not revealed significant differences in either contraction times (Figure S5B-C) or relaxation times (Figure S5D) for the *LMNA* mutant iPSC-CMs compared to either of the healthy controls.

To determine if the nuclear size and shape defects could be attributed to altered actin and microtubules organization or cytoskeletal forces resulting from these networks, we targeted these networks in a representative healthy control iPSC-CM cell line, WT2, and the R249Q iPSC-CMs, representing the most severe nuclear damage. Cells were treated either with drugs to depolymerize (Latrunculin B) or inhibit contractility (Blebbistatin) of the actin network, or to stabilize (Paclitaxel) or depolymerize (Nocodazole) microtubules. We confirmed the intended effect of each drug on actin and microtubules organization, respectively (Figure 5A,B). In addition, we immunofluorescently labeled Lamin B1 to quantify nuclear area and circularity index following drug treatment or vehicle control treatment. When normalized to the vehicle (DMSO) control, only the actomyosin contractility inhibitor, Blebbistatin, and the microtubule-stabilizing drug, Paclitaxel, induced a slight increase in nuclear area in R249Q iPSC-CMs compared to DMSO controls (Figure 5C and E, respectively). Treatment with Nocodazole, which depolymerizes microtubules, slightly increased R249Q iPSC-CM nuclear circularity compared to DMSO controls, although this effect was small (<5%) and only reached statistical significance when comparing results to vehicle controls (Figure 5F), and not when comparing absolute circularity index values across all groups (Figure S6). None of the drug treatments induced changes in WT2 iPSC-CM nuclear area or circularity index (Figure 5D and F, respectively) compared to DMSO controls. Taken together, these results suggest that actin contractility plays only a small role in the increased nuclear size in R249Q iPSC-CMs. In contrast to a recent report linking microtubule mediated forces to nuclear defects in cardiac laminopathies (Leong *et al*., 2023), we observed only a small effect when disrupting the microtubule network, which may be explained at least in part by the less mature cytoskeletal organization in iPSC-CMs compared to adult cardiaomyocytes. Taken together, our results suggest that the severe changes in nuclear shape and nuclear envelope rupture are predominantly due to intrinsic nuclear defects, such as altered Lamin A/C protein expression and assembly, rather than due to differences in cytoskeletal forces acting on the nuclei.

**Figure 5.**
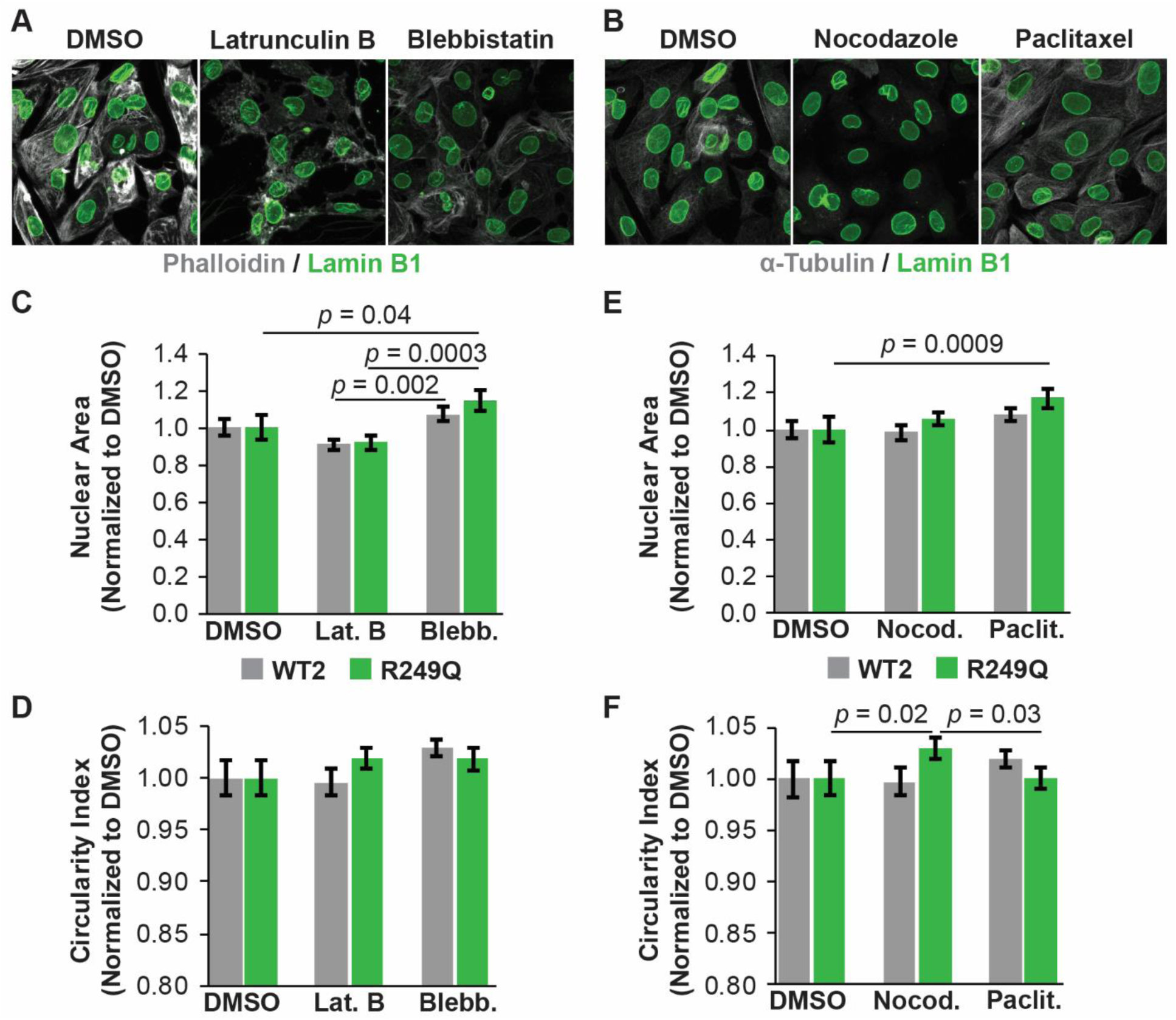
The role of the cytoskeleton in R249Q iPSC-CM nuclear damage and fragility. (A) Representative Phalloidin and Lamin B1 images of R249Q iPSC-CMs treated with either a vehicle (DMSO) control, Latrunculin B to depolymerize the actin cytoskeleton, as evidenced by fragmentation of actin, or Blebbistatin to inhibit actin contractility. Scale bar = 50 µm. (B) Representative α-Tubulin and Lamin B1 images of R249Q iPSC-CMs treated with either a vehicle (DMSO) control, Nocodazole to depolymerize microtubules, as evidenced by the ablation of microtubule structure, or Paclitaxel to stabilize microtubules. Scale bar = 50 µm. (C) Actin-targeting drugs Latrunculin B and Blebbistatin did not impact nuclear area of WT2 iPSC-CMs, while only Blebbistatin significantly increased nuclear area in R249Q iPSC-CM compared to the vehicle (DMSO) control. (D) Actin-targeting drugs Latrunculin B and Blebbistatin did not induce changes in circularity index for either WT2 or R249Q iPSC-CMs compared to the vehicle control. (E) Microtubule-targeting drugs Nocodazole and Paclitaxel did not impact nuclear area of WT2 iPSC-CMs, while only Paclitaxel significantly increased nuclear area in R249Q iPSC-CM compared to the vehicle (DMSO) control. (F) Microtubule-targeting drugs Nocodazole and Paclitaxel did not alter nuclear area of WT2 iPSC-CMs, while only Nocodazole significantly increased circularity index in R249Q iPSC-CM compared to the vehicle (DMSO) control. Data presented as mean ± SEM. *N* ≥ 44 nuclei per group over two to three independent experiments for nuclear area and circularity index.

### LMNA R249Q-iPSC-CMs have reduced nuclear stiffness

Since cytoskeletal disruption had only a minor effect on nuclear shape, we investigated the effect of *LMNA* mutations on nuclear stiffness. Lamin A/C are a major determinant of nuclear stiffness (Lammerding *et al*., 2004, 2006; Swift *et al*., 2013; Stephens *et al*., 2017), and *LMNA*-mutations can cause changes in nuclear stability that lead to nuclear damage (Sullivan *et al*., 1999; Nikolova *et al*., 2004; Gupta *et al*., 2010; De Vos *et al*., 2011; Zwerger *et al*., 2013; Cho *et al*., 2019; Earle *et al*., 2020). To determine whether the high degree of nuclear defects exhibited by R249Q mutant nuclei could be related to altered nuclear mechanics, we measured nuclear stiffness of R249Q mutant iPSC-CMs and healthy controls with a nanoindenter using a spherical probe tip (Figure 4E). We confirmed that nuclei in iPSC-CMs are close to the cell surface with little to no cytoskeleton over the top of nuclei (Figure S7A). Thus, when probing the nucleus in intact cells, the resulting force-indentation curves are expected to primarily reflect the mechanical properties of the cell nucleus. The R249Q iPSC-CMs had significantly reduced nuclear stiffness (Figure 4F), which was also reflected by the fact that at the same indentation force, the R249Q mutant iPSC-CM nuclei had a significantly increased indentation depth compared to healthy control iPSC-CMs (Figure S7B). Interestingly, although R249Q mutant iPSC-CMs had increased nuclear cross-sectional areas compared to healthy control iPSC-CMs (Figure S7C), consistent with our earlier measurements (Figure 2D), we did not find a significant correlation between nuclear cross-sectional area and nuclear stiffness across individual cells for either cell line (Figure S7D), likely due to the large variability in nuclear cross-sectional area between individual cells.

Since a previous study observed increased nuclear envelope rupture in Lamin A/C-depleted iPSC-CMs at low indentation forces (10-30 nN) (Xia *et al*., 2018), we tested whether nanoindentation also increased nuclear envelope rupture in our experimental system. However, neither healthy control nor R249Q iPSC-CMs exhibited nuclear envelope ruptures during indentation, neither at the 40 nN load used for nuclear stiffness measurements, nor when increasing the indentation force up to 200 nN with a range of probe sizes (3, 10, or 50 μm radius). The lack of nuclear envelope rupture in our experiments is likely explained by the larger probe size required for our nanoindentation system compared to the probes used in the previous study (<0.1 µm diameter), as previous studies found that high nuclear envelope curvature induced with sharp AFM probes promote local nuclear envelope defects and nuclear envelope rupture (Xia *et al*., 2018; Pfeifer *et al*., 2022).

### Reduced Lamin A/C expression can partially explain R249Q nuclear damage

Since the mutant iPSC-CM lines have a heterogeneous genetic background, we wanted to determine if the defects in nuclear shape and stability observed were due mutation of *LMNA* or genetic background differences. To test for this, we assessed the effects of shRNA-mediated Lamin A/C depletion (shLMNA) in healthy control iPSC-CMs on nuclear morphology and nuclear envelope rupture and compared the results with those obtained for a non-target shRNA (shNT) in isogenic control cells. Depletion of Lamin A/C was confirmed by immunofluorescence labeling. Due to limited transduction efficiency in iPSC-CMs, only about 50% of the shLMNA-modified cells exhibited loss of Lamin A/C (‘shLMNA-KD’ cells) at 7 days post transfection, with the remainder showing near normal Lamin A/C levels (‘shLMNA-no KD’ cells) (Figure 6A; Figure S8A-B). The shLMNA-KD iPSC-CMs had significantly increased nuclear area and volume compared to shLMNA-no KD and shNT controls (Figure 6C-D), consistent with the increase in nuclear area and volume observed in the R249Q cells (Figure 3D-E). Surprisingly, we did not detect any changes in circularity index between shLMNA-KD nuclei and either shNT or shLMNA-no KD nuclei (Figure 6B). These findings suggest that reduced Lamin A/C levels directly influence nuclear size in iPSC-CM nuclei, but that other factors may contribute to abnormal nuclear shape in the *LMNA* mutant iPSC-CMs. However, it is also possible that prolonged loss of Lamin A/C is required to cause more substantial defects in nuclear shape, or that the timing of Lamin A/C loss during cardiac differentiation of iPSC-CMs influences nuclear shape, which is further supported by a recently discovered role of microtubule mediated forces on nuclear position and defects in Lamin A/C deficient cardiomyocytes (Leong *et al*., 2023).

**Figure 6.**
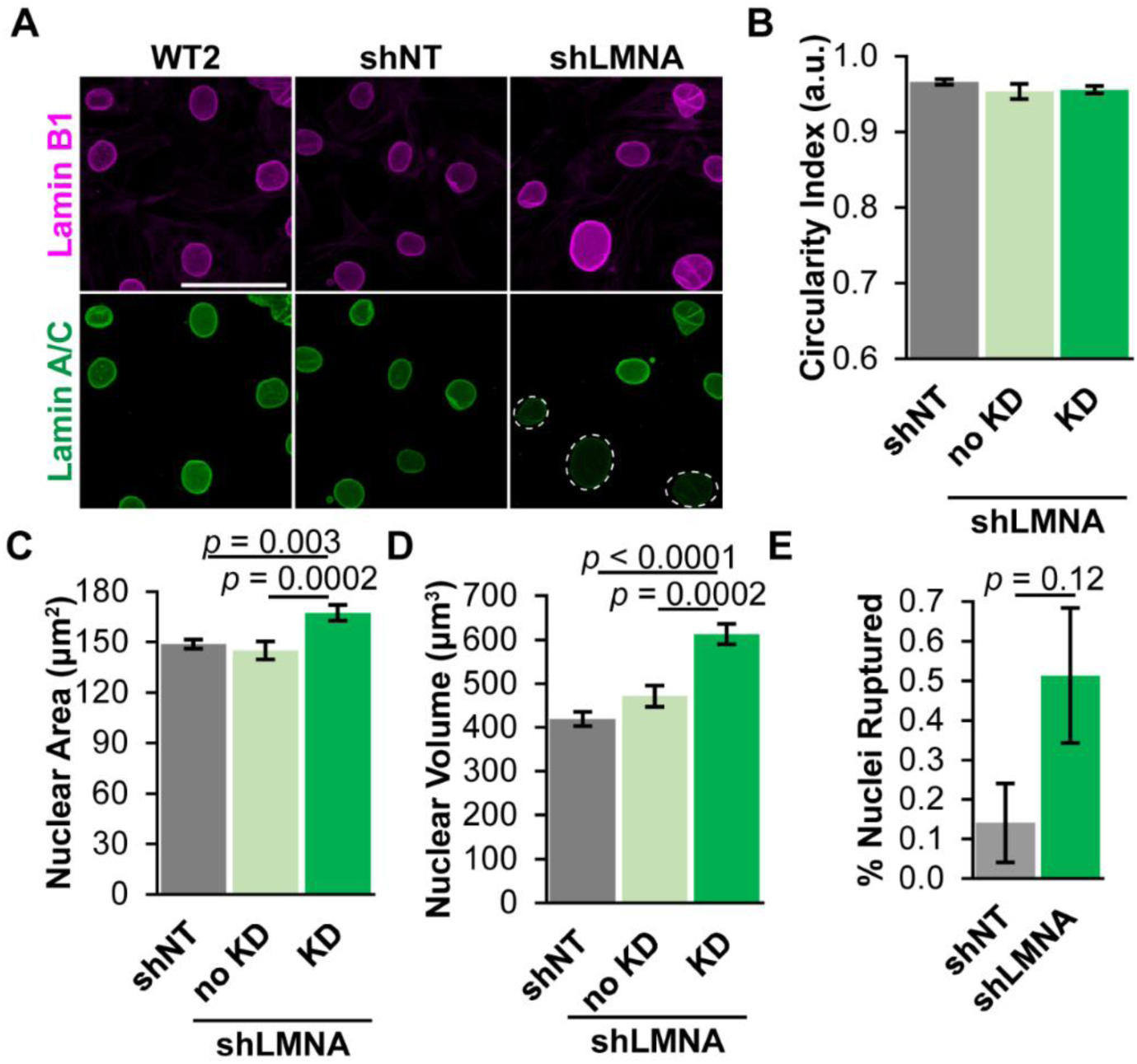
Depletion of Lamin A/C in healthy control iPSC-CMs causes increased nuclear area, volume, and nuclear envelope ruptures. (A) Representative immunofluorescence images for Lamin B1 and Lamin A/C in healthy control (WT) iPSC-CMs either modified with shRNA targeting *LMNA* (shLMNA) or a non-target control shRNA (shNT). White dotted circles mark shLMNA treated nuclei with visible Lamin A/C depletion. Scale bar = 50 µm. (B) shLMNA nuclei with depletion of Lamin A/C (knockdown; KD shLMNA) have no change in circularity index compared to both non-target (shNT) control and non-depleted shLMNA nuclei (no KD). (C) KD shLMNA nuclei had significantly increased nuclear area compared to both non-target (shNT) control and non-knockdown shLMNA nuclei. (D) KD shLMNA nuclei had significantly increased nuclear volume compared to both non-target (shNT) control and non-knockdown shLMNA nuclei. (E) Unselected populations of shLMNA treated iPSC-CMs had a trend towards increased nuclear envelope rupture compared to shNT treated controls. Due to the experimental protocol using live cells, we could not distinguish cells with successful Lamin A/C depletion from those without Lamin A/C depletion in the shLMNA treated cells. Data presented as mean ± SEM. *N* ≥ 83 nuclei per group for nuclear shape and area, *N* ≥ 47 nuclei per group for nuclear volume, *N* > 1,415 nuclei per group for nuclear envelope rupture.

Time-lapse experiments with shLMNA- and shNT-modified iPSC-CMs expressing the NLS-GFP nuclear envelope rupture reporter suggested that Lamin A/C depletion increases nuclear envelope rupture (Figure 6E), although the difference between the shLMNA iPSC-CMs and shNT controls was only trending towards statistical significance (*p* = 0.12). This lack of statistical significance was likely due to the limited transduction efficiency, in which only ∼50% of the shLMNA cells had reduced levels of Lamin A/C (Figure S8B). Overall, these data are consistent with a previous study reporting increased nuclear envelope rupture in Lamin A/C depleted cardiomyocytes (Cho *et al*., 2019), and suggest that reduced Lamin A/C levels or functional loss of Lamin A/C could be at least partially responsible for the increase in nuclear envelope ruptures in the R249Q mutant iPSC-CMs. On the other hand, the fraction of shLMNA iPSC-CMs exhibiting nuclear envelope rupture was substantially lower than in the R249Q mutant cells, even when accounting for the reduced transduction efficiency, suggesting that additional mechanism contribute to the increased incidence of nuclear envelope rupture in the R249Q mutant iPSC-CMs.

### LMNA mutant iPSC-CMs exhibit reduced localization of lamins to the nuclear envelope

*LMNA* mutations may alter Lamin A/C self-assembly or lamin phosphorylation, resulting in improper assembly of the lamina and mislocalization of Lamin A/C from the nuclear envelope (Cenni *et al*., 2005; Wiesel *et al*., 2008; Zwerger *et al*., 2015; Bertrand *et al*., 2020). Since defects in lamin assembly could impair the resistance of the nucleus to mechanical forces exerted by the cytoskeleton, resulting in nuclear defects and nuclear envelope rupture (De Vos *et al*., 2011; Zwerger *et al*., 2013; Denais *et al*., 2016), we examined the intranuclear distribution of Lamin A/C and Lamin B1 in the *LMNA* mutant and healthy control iPSC-CMs by immunofluorescence labeling. Both the R249Q and the L35P mutant iPSC-CMs frequently had reduced Lamin A/C fluorescence at the nuclear periphery (Figure 7A). For a more quantitative analysis, we measured the immunofluorescence intensity profiles of Lamin A/C (Figure 7B) and Lamin B1 (Figure S9) along a line across the central z-plane of nuclei. Lamin fluorescence intensity profiles were normalized to the nuclear diameter and to the area under the curve to account for variations in nuclear size and Lamin A/C levels (Figure 7C). Whereas healthy control iPSC-CMs exhibited Lamin A/C fluorescence intensity profiles with large peaks at the nuclear periphery, indicating that Lamin A/C is predominantly found at the nuclear lamina in these cells, the R249Q, L35P, and, to a lesser extent, G449V mutant iPSC-CMs had significantly decreased Lamin A/C at the nuclear periphery and increased Lamin A/C in the nucleoplasm (Figure 7C-D).

**Figure 7.**
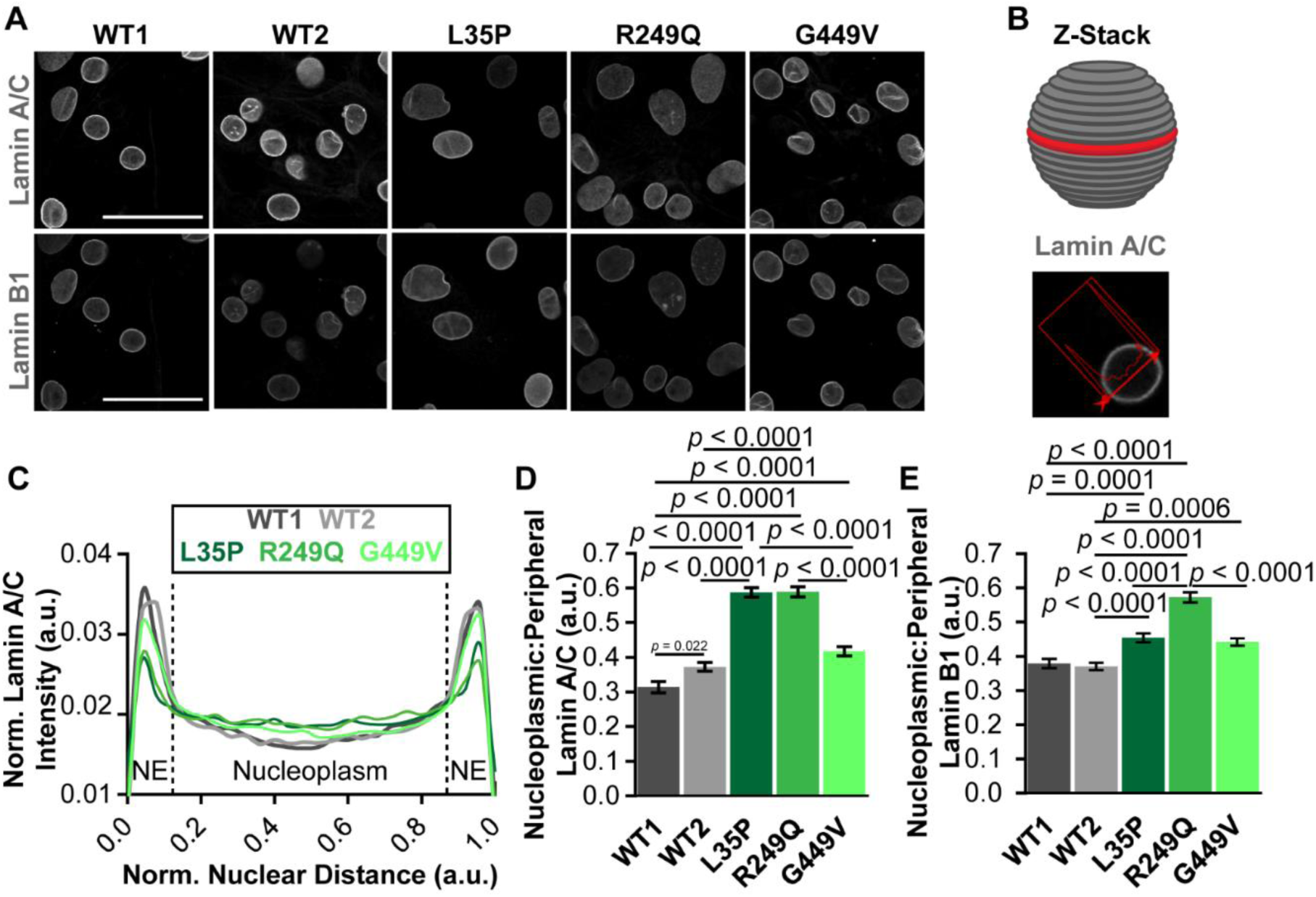
*LMNA*-mutant iPSC-CMs have mislocalization of Lamin A/C and Lamin B1 from the nuclear envelope. (A) Confocal immunofluorescence images of cross sections through the center of nuclei show that L35P and R249Q iPSC-CM nuclei have reduced fluorescence intensity of Lamin A/C at the nuclear periphery, and R249Q nuclei additionally have reduced fluorescence intensity of Lamin B1 at the nuclear periphery. Scale bars = 50 µm. (B) Schematic of the lamin localization profile analysis. Lamin A/C fluorescence intensity profiles were taken at the central z-plane of nuclei from confocal image slices. (C) A plot of the normalized Lamin A/C fluorescence intensity profile shows that L35P and R249Q nuclei have reduced fluorescence intensity peaks at the nuclear periphery (nuclear envelope; NE) and increased fluorescence intensity in the nucleoplasm. (D) L35P, R249Q, and G449V iPSC-CMs had a significantly increased ratio of nucleoplasmic to peripheral Lamin A/C, indicating lamin mislocalization away from the nuclear envelope. (E) L35P, R249Q, and G449V-iPSC-CMs have a significantly increased ratio of nucleoplasmic to peripheral Lamin B1, with R249Q nuclei having a particularly increased ratio, indicating lamin mislocalization away from the nuclear envelope. Data presented as mean ± SEM. *N* > 60 nuclei per group.

To understand the relative contributions of assembled versus soluble Lamin A/C to lamin mislocalization, we washed out soluble nuclear proteins in semi-permeabilized nuclei of healthy control and R249Q mutant iPSC-CMs, as a representative cell line. We imaged the remaining non-soluble Lamin A/C (Figure S10A) and quantified the un-normalized Lamin A/C fluorescence intensity in the nucleoplasm and at the nuclear envelope to better capture the differences in protein levels at each location (Figure S10B). Both R249Q mutant and healthy control iPSC-CMs had a similar pool of nucleoplasmic Lamin A/C that decreased with washout (Figure S10B), indicating that much of the nucleoplasmic Lamin A/C was soluble and that R249Q mutant iPSC-CMs maintain a similar pool of nucleoplasmic Lamin A/C to healthy controls. However, the R249Q mutant iPSC-CMs had significantly decreased Lamin A/C at the nuclear envelope compared to healthy controls, both in non-washout and washout conditions (Figure S10B), suggesting that the R249Q mutation results in reduced Lamin A/C assembly at the NE. Lamin A/C fluorescence intensity profiles for non-washout (Figure S10C) and washout (Figure S10D) iPSC-CMs provide additional visual confirmation of these results. Together, these results suggest that the changes in Lamin A/C localization can be attributed to reduced levels of Lamin A/C assembled at the NE.

Lamin A/C and Lamin B1 form separate but interacting protein meshworks at the nuclear periphery (Shimi *et al*., 2015; Xie *et al*., 2016), and mutation or depletion of either protein can disrupt the structural organization of the other (Vigouroux *et al*., 2001; Muchir *et al*., 2004; Guo *et al*., 2014; Shimi *et al*., 2015; Steele-Stallard *et al*., 2018). Thus, we hypothesized that perturbed Lamin A/C assembly could also impact the organization of Lamin B1 in the *LMNA* mutant iPSC-CMs. Indeed, all three *LMNA* mutant iPSC-CM lines exhibited an increased nucleoplasmic to peripheral ratio of Lamin B1 compared to healthy control iPSC-CMs (Figure 7E), with R249Q iPSC-CMs exhibiting the most severe defects in Lamin B1 intranuclear distribution, and L35P and G449V iPSC-CMs displaying milder abnormalities. Collectively, these data indicate that mislocalization of Lamin A/C from the nuclear envelope can alter Lamin B1 distribution. Furthermore, our findings suggest that the particularly severe defects in nuclear morphology and stability in the R249Q mutant iPSC-CMs could arise in part from the mislocalization of Lamin B1 from nuclear periphery, given the critical role of B-type lamins in nuclear stability and preventing nuclear envelope rupture (Lammerding *et al*., 2006; Hatch *et al*., 2013; Chen *et al*., 2018, 2019; Pfeifer *et al*., 2022).

### Lamin A/C phosphorylation is not altered in LMNA mutant iPSC-CMs

An altered ratio of nucleoplasmic to peripheral Lamin A/C could be due to a reduced ability of mutant Lamin A/C to self-assemble into higher order filaments (Wiesel *et al*., 2008; Zwerger *et al*., 2013; Bertrand *et al*., 2020) or an increase in Lamin A/C phosphorylation, which inhibits Lamin A/C’s ability to form dimers and assemble at the nuclear periphery (Swift *et al*., 2013; Buxboim *et al*., 2014). To test whether increased phosphorylation of Lamin A/C was responsible for reduced Lamin A/C localization to the NE, we immunofluorescently labeled iPSC-CMs for Lamin A/C phosphorylated at Serine 22 (p-Lamin A/C) and for total Lamin A/C (Figure S11A). We did not detect a significant increase in the ratio of phosphorylated Lamin A/C to total Lamin A/C in the *LMNA* mutant iPSC-CMs (Figure S11B), indicating that increased Lamin A/C phosphorylation is not responsible for reduced peripheral Lamin A/C in these cells. Thus, these data suggest that altered self-assembly of Lamin A/C into higher order filaments at the nuclear envelope is likely responsible for lamin mislocalization in *LMNA*-mutant iPSC-CMs.

### Nuclear abnormalities correlate with lamin mislocalization from the nuclear envelope

Our data suggest that *LMNA* mutant iPSC-CMs have, to a varying degree, defects in Lamin A/C and Lamin B1 localization to the nuclear periphery and that the R249Q mutant iPSC-CMs show particularly severe defects in both Lamin A/C and Lamin B1 organization, an observation which may explain the increased susceptibility of these cells to nuclear damage. To determine the degree to which mislocalization of lamins can explain nuclear size and shape defects in different cell lines, we computed a “lamin mislocalization index” from the average of the nucleoplasmic to peripheral Lamin A/C and Lamin B1 ratios for each cell line and correlated this index with a normalized “nuclear defects score,” computed from the geometric mean of nuclear circularity index, area, and volume (Figure 8). Intriguingly, the nuclear defects score across the panel of iPSC-CMs strongly correlated with the lamin mislocalization index (R^2^ = 0.96) with a significantly positive slope (*p* = 0.004), indicating that the extent of impaired Lamin A/C and Lamin B1 assembly at the nuclear envelope can explain much of the varying degrees of nuclear damage severities across multiple cell lines. Although other mechanisms, such as cytoskeletal forces exerted on the nucleus or reduced Lamin A/C protein expression may account for some of the remaining portion of the variability, our findings point to an important role of Lamin A/C—and also Lamin B1—mislocalization from the nuclear envelope to the cytoplasm in predicting the effect of specific *LMNA* mutations on nuclear damage in iPSC-CMs.

**Figure 8.**
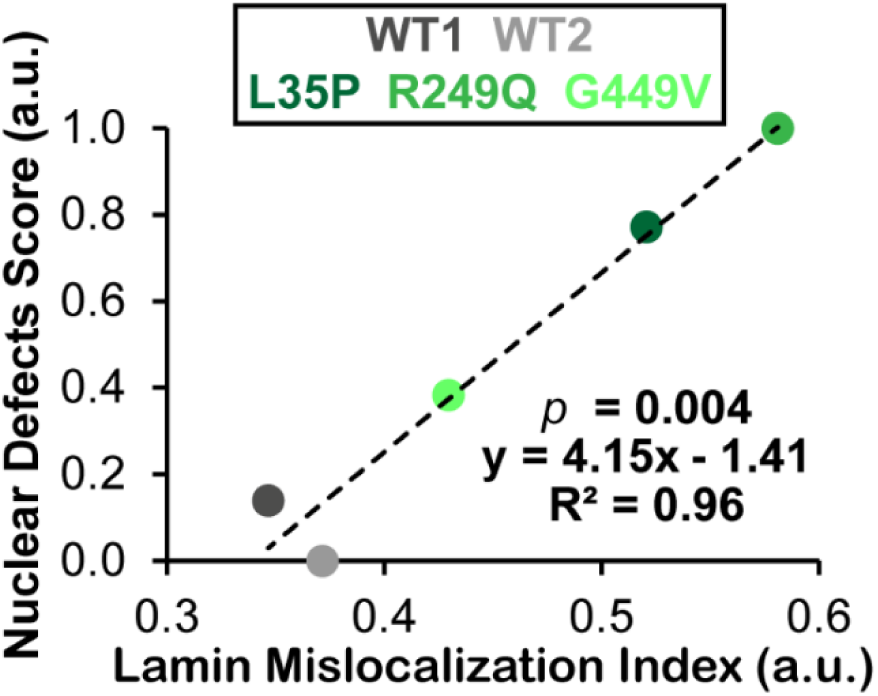
Lamin mislocalization correlates to nuclear damage in *LMNA*-mutant iPSC-CMs. Lamin mislocalization index (defined as the average of Lamin A/C and Lamin B1 nucleoplasmic to peripheral intensity ratios) shows a significant positive correlation to the nuclear defects score (defined as the normalized geometric mean of nuclear circularity index, area, and volume) for the five iPSC-CM cell lines used in this study.

## Discussion

The mechanisms by which different *LMNA* mutations cause an array of disease phenotypes and severities has long remained an open question. Here, using three *LMNA* mutant iPSC lines associated with *LMNA*-DCM, we demonstrate that the degree of impaired localization of Lamin A/C and Lamin B1 to the nuclear envelope strongly correlates with the severity of nuclear damage. We propose a mechanism in which *LMNA* mutations cause defective assembly of Lamin A/C, resulting in reduced Lamin A/C levels at the nuclear periphery, which, in turn, perturbs Lamin B1 localization to the nuclear periphery. Collectively, this reduced lamin assembly at the nuclear envelope leads to impaired nuclear stability, resulting in defects in nuclear shape and size and increased nuclear fragility and nuclear envelope rupture (Figure 9). In contrast, altered cytoskeletal forces exerted on the nucleus appear to have only a small role in causing nuclear damage in the *LMNA* mutant iPSC-CMs, although they may play a more prominent role in adult cardiomyocytes, which generate substantially higher forces, or over longer periods of time.

**Figure 9.**
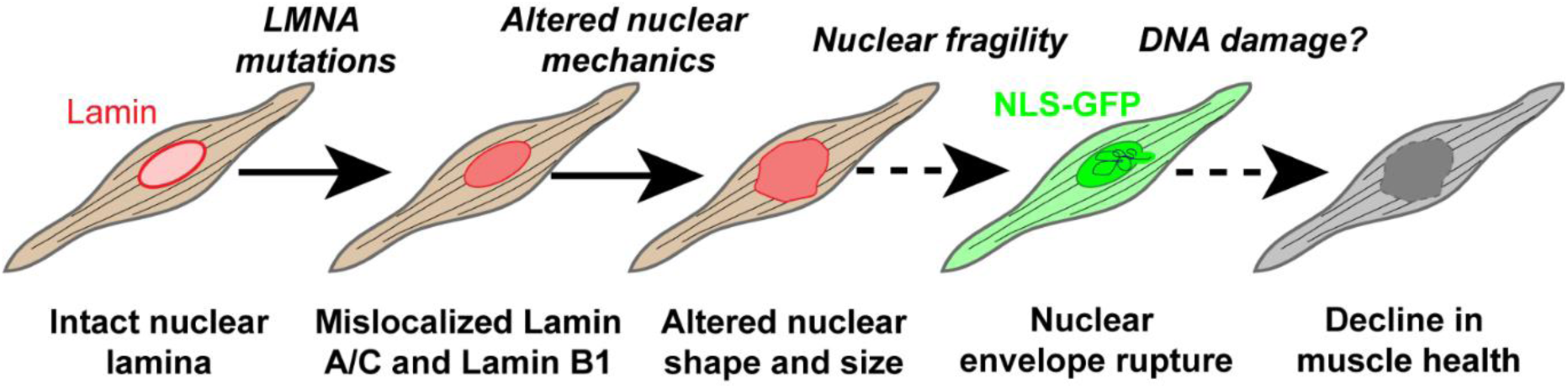
Proposed mechanism of nuclear damage in *LMNA*-mutant iPSC-CMs. Reduced Lamin A/C levels and/or defective assembly of Lamin A/C causes Lamin A/C mislocalization from the nuclear envelope. This mislocalization alters nuclear mechanics, particularly reducing nuclear stiffness and stability, leaving nuclei susceptible to cytoskeletal forces exerted on the nucleus in cardiomyocytes. These forces in turn cause abnormal nuclear size and shape and increased nuclear envelope rupture. Nuclear envelope rupture, and potentially other nuclear damage, can cause a decline in muscle health, potentially through mechanisms such as DNA damage or increased apoptosis.

Similar to iPSCs from healthy controls, *LMNA* mutant iPSCs have low expression of Lamin A/C and lack nuclear shape abnormalities. Upon differentiation into cardiomyocytes, Lamin A/C expression is upregulated, and *LMNA* mutant iPSC-CMs display varying degrees of nuclear abnormalities in the form of altered nuclear shape and/or increased nuclear size compared to healthy controls. R249Q mutant iPSC-CMs had the most severe nuclear size and shape defects and exhibited significantly more nuclear envelope ruptures than the other iPSC-CMs, likely due to their reduced nuclear stiffness and stability. Healthy control iPSC-CMs depleted of Lamin A/C showed similar defects in nuclear morphology and increased nuclear envelope rupture, as in the *LMNA* mutant iPSC-CMs, particularly the R249Q cells, confirming our observations in an isogenic background.

Intriguingly, R249Q mutant iPSC-CMs exhibited higher levels of nuclear envelope rupture than the Lamin A/C-depleted iPSC-CMs. Furthermore, both healthy control and *LMNA*-mutant iPSC-CMs exhibited substantial variation in overall levels of Lamin A/C, although only the L35P and R249Q iPSC-CMs exhibited significantly decreased Lamin A/C expression compared to both or one healthy control, respectively. These reduced Lamin A/C levels may be indicative of partial *LMNA* haploinsufficiency, caused by altered translation and/or turnover of the mutant Lamin A/C, which has been previously shown to cause *LMNA*-DCM (Cattin *et al*., 2013). Depletion of Lamin A/C in healthy control iPSC-CMs demonstrated that nuclear size and to some extent nuclear envelope rupture are mediated by Lamin A/C levels, and therefore a degree of *LMNA* haploinsufficiency in R249Q iPSC-CMs could in part explain the increase of nuclear size and nuclear envelope ruptures. In support of this idea, expression of a *Drosophila* Lamin C mutant corresponding to the human R249Q mutation caused dominant effects on nuclear structure, including alterations in the organization of A- and B-type lamins (Hinz *et al*., 2021). Furthermore, *in silico* analysis of the Lamin A/C R249Q dimer predicted an abnormal arched conformation in the alpha helical rod domain of the mutant Lamin A/C model, which could be responsible for altered lamina assembly (Hinz *et al*., 2021).

Pointing to a potential fundamental principle, all three *LMNA* mutant iPSC-CM lines exhibited Lamin A/C and Lamin B1 mislocalization from the nuclear periphery compared to healthy controls. This is likely due to defective assembly of Lamin A/C at the nuclear lamina, as the *LMNA* mutant cells still showed reduced Lamin A/C levels at the nuclear envelope after removing soluble lamins (Figure S9). These results are in line with previous studies showing that *LMNA* mutations may result in Lamin A/C mislocalization from the nuclear envelope (Wiesel *et al*., 2008; Zwerger *et al*., 2013; Steele-Stallard *et al*., 2018; Bertrand *et al*., 2020), and that disrupting Lamin A/C assembly at the nuclear periphery impairs nuclear stability (Zwerger *et al*., 2013). It is now well recognized that Lamin A/C and Lamin B1 form independent and interacting meshworks at the nuclear periphery, and that depletion of one filament system disrupts the structure of the other (Vigouroux et al., 2001; Muchir et al., 2004; Guo et al., 2014; Shimi et al., 2015; Steele-Stallard et al., 2018). However, little is understood about the consequences of altered Lamin A/C assembly on Lamin B1 organization in laminopathies. In Drosophila laminopathy models, muscle-specific expression of specific mutant A-type lamins caused mislocalization of B-type lamins (Dialynas *et al*., 2012). Our results demonstrate in iPSC-CMs, *LMNA* mutations can displace Lamin B1 from the NE, which is expected to make cells more susceptible to nuclear envelope rupture (Lammerding *et al*., 2006; Hatch *et al*., 2013; Denais *et al*., 2016).

Whereas previous studies have shown that Lamin A/C mislocalization correlates with defective nuclear shape (Wiesel *et al*., 2008; Zwerger *et al*., 2013; Steele-Stallard *et al*., 2018; Bertrand *et al*., 2020) and altered nuclear mechanics (Lammerding *et al*., 2004; Zwerger *et al*., 2015), our results point to an intriguing discovery that the ‘lamin mislocalization index,’ describing defects in both Lamin A/C and Lamin B1 assembly at the NE, can explain >95% of the variability in nuclear defects in different *LMNA* mutations and healthy control cell lines. As noted previously, Lamin A/C haploinsufficiency, and to a smaller extent the exertion of cytoskeletal forces on the nucleus, could contribute to the nuclear morphological defects, which could explain some of the remaining 5% of the variability in nuclear morphological defects. Interestingly, at the level of individual nuclei, this correlation between lamin mislocalization and degree of nuclear abnormalities is roughly maintained among healthy control iPSC-CM nuclei, but not R249Q nuclei (Figure S12), suggesting that the increase in nuclear defects observed in R249Q mutant iPSC-CMs cannot be solely explained by an increase in lamin mislocalization from the nuclear periphery. This effect could be due to a threshold for the amount of lamin assembled at the NE, below which nuclei are increasingly susceptible to nuclear abnormalities. R249Q mutant iPSC-CMs generally have a high degree of both Lamin A/C and Lamin B1 mislocalization from the nuclear envelope and significantly decreased nuclear stiffness, which could indicate that their nuclei are below this threshold for peripheral lamin assembly and thus are more susceptible to deformation and nuclear envelope rupture.

The loss of nuclear mechanical strength and nuclear envelope integrity due to the loss of Lamin A/C and/or Lamin B1 at the nuclear envelope can have severe consequences on cellular health and survival (Denais et al., 2016; Chen et al., 2018, 2019; Earle et al., 2019; Shah et al., 2021), particularly through susceptibility of the nucleus to damage from intracellular forces (Hatch and Hetzer, 2016; Cho et al., 2019; Earle et al., 2019b; Shah et al., 2021). One notable observation of our study, however, is that although the degree of lamin mislocalization is a reasonable predictor of the effect of the *LMNA* mutation on nuclear mechanics and stability, it is not necessarily a good predictor of patient disease phenotype. Of the three *LMNA* mutations used in this study, the patient with the L35P mutation had the most severe phenotype, presenting both skeletal muscle and cardiac phenotypes during childhood and living only to age 15. Conversely, the patient presenting the R249Q mutation, whose iPSC-CMs had more severe disruptions to nuclear mechanics than L35P iPSC-CMs, developed early onset muscular dystrophy in childhood, but only developed severe cardiac dysfunction later in life. This disparity between patient and cellular severity is not completely surprising, as additional defects could be due to altered gene expression and/or biochemical signaling that occur during development, which will affect disease onset and progression. As such, altered Lamin A/C and Lamin B1 assembly at the nuclear envelope can cause changes in chromatin organization and gene expression (Bertero *et al*., 2019; Cheedipudi *et al*., 2019; Kim *et al*., 2019), including nuclear mechanotransduction and associated signaling (Kirby and Lammerding, 2018; Maurer and Lammerding, 2019; Donnaloja *et al*., 2020). Thus, loss of assembled Lamin A/C and/or Lamin B1 at the nuclear periphery may impair mechanically-induced gene expression (Maurer and Lammerding, 2019; Donnaloja *et al*., 2020).

### Limitations of the current study

We recognize that this study has several limitations. First, our study is limited by the lack of isogenic cell lines available for comparison, as the heterogeneous genetic background of the iPSC lines used here could potentially conflate differences in genetic backgrounds with differing *LMNA*-DCM phenotypes. Unfortunately, despite extensive efforts, we were unable to generate isogenic controls for the various *LMNA* mutant iPSC lines. Thus, although we observed only small differences in phenotype between the two healthy control iPSC-CM lines in this study and we used shRNA-mediated depletion to recapitulate certain aspects of *LMNA*-DCM nuclear morphological defects and nuclear envelope rupture in an isogenic background, we cannot rule out that additional factors, including genetic variations in modifier genes or other structural proteins, could contribute to the nuclear morphologies observed in this study.

In addition, the iPSC-CMs used in this study are immature compared to adult cardiomyocytes, lacking the highly organized cytoskeletal organization and other features (Ahmed *et al*., 2020; Wu *et al*., 2021). Although we did not observe any differences in contractility between *LMNA* mutant iPSC-CMs and healthy controls and thus ruled out a role for altered contractility in nuclear morphological defects, our results cannot exclude the possibility that other metrics not included in our study, such as calcium handling and contractility forces, may reveal contractility defects in the *LMNA* mutant iPSC-CMs used. Furthermore, *LMNA* mutant mature cardiomyocytes may exhibit functional alterations not apparent in the immature iPSC-CMs, as suggested by other studies (Bertero *et al*., 2019; Mehrabi *et al*., 2021; Miura *et al*., 2022; Wang *et al*., 2022b).

Finally, several additional factors not investigated thoroughly in this study may contribute to remainder of the variability in nuclear defects. Our results modifying actin and microtubule contractility or assembly yielded only very modest changes in R249Q iPSC-CM nuclear shape and size in response to some drug treatments, but consistent trends did not emerged. Therefore, although we postulate that lamin expression and localization are the predominant factors responsible for the severe nuclear damage in the R249Q iPSC-CMs, we did not investigate the extent to which actin contractility versus microtubule associated forces cause nuclear damage, their respective roles in in nuclear envelope rupture (Cho *et al*., 2019; Pfeifer *et al*., 2022; Leong *et al*., 2023), or other changes to the cell or nuclear structure, such as altered expression or organization or nuclear envelope proteins or disrupted nucleo-cytoskeletal coupling (Lammerding et al., 2004; Hale et al., 2008; Chen et al., 2012; Arsenovic et al., 2016). Furthermore, in our investigation, iPSC-CMs were neither uniformly aligned on a substrate nor electrically paced, thus resulting in disorganized cellular structure, which likely increased cell-to-cell variability in the contractility measurements. Thus, we cannot exclude that the *LMNA* mutations studied here could display effects on myocyte contractility in other settings. Finally, an additional possible factor for lamin mislocalization that we did not investigate in this study is the role of the cell cycle, which is known to affect intranuclear localization of Lamin A/C (Boban *et al*., 2010). However, iPSC-CMs are largely post-mitotic, and division of iPSC-CM was extraordinarily rare in our live cell imaging experiments. Thus, we believe that cell cycle-dependent effects are a relatively negligible factor in the degree of lamin mislocalization observed in *LMNA* mutant iPSC-CMs.

### Conclusions

Our results point to lamin mislocalization from the nuclear envelope as a common potential pathogenic mechanism across *LMNA*-DCM mutations, which determines the degree of nuclear mechanical damage in cardiomyocytes. Moreover, altered lamin assembly at the nuclear envelope may present a potential link between the proposed laminopathy pathogenic mechanisms of impaired nuclear stability and altered gene expression driving *LMNA* mutant skeletal muscle and cardiac laminopathies (Maurer and Lammerding, 2019; Donnaloja *et al*., 2020). Future studies should be aimed at determining the molecular mechanisms by which nuclear damage causes cardiac dysfunction in *LMNA*-DCM, including obtaining a better understanding of changes in mechanotransduction signaling and gene expression. Additionally, our work and others showing that Lamin A/C assembly has implications in the degree of nuclear damage and altered nuclear mechanics (Wiesel *et al*., 2008; Zwerger *et al*., 2013; Bertrand *et al*., 2020) suggest that by understanding the mutation-specific degree of defective lamin assembly, we may ultimately be able to identify patient mutations that would most benefit from therapeutics targeting the reduction of force transmission to the nucleus (Cho *et al*., 2019; Earle *et al*., 2020; Chai *et al*., 2021). Although additional mechanisms may drive *LMNA*-DCM, such therapeutics hold promise to improve cellular health and survival (Cho *et al*., 2019; Earle *et al*., 2020; Chai *et al*., 2021).

## Acknowledgements

The authors thank the Cure Muscular Dystrophy Foundation for the *LMNA* L35P iPSCs; Hanna Gimse for analyzing the nuclear volume in shRNA experiments; the Cornell Institute of Biotechnology for performing library preparation and Illumina sequencing experiments; the Cornell Statistical Consulting Unit for developing the regressions performed in R; Dr. Kehan Zhang and Dr. Christopher Chen for hosting M.W. to learn iPSC culture and cardiac differentiation protocols; and Dr. Kathleen Xie for the nuclear semi-permeabilization protocol. This work was supported by awards from the National Institutes of Health (R01 HL082792 to J.L., R01 HL128075 to E.M.M., and F30 HL142187 to A.M.G.); the National Science Foundation (CBET 1715606 and URoL-2022048 to J.L.; Graduate Research Fellowships 2016229710 to M.W.), the American Heart Association (20PRE35080179 to M.W.), the Muscular Dystrophy Association (MDA603238 to T.J.K), and the Volkswagen Foundation (A130142 to J.L.).

## Notes

### Competing Interest Statement

The authors have declared no competing interest.

### Summary of Updates

We have added a new Supplementary Figure (S6) showing the non-normalized data for the nuclear circularity index in cells treated with Nocodazole or vehicle control. We have revised Figure 5 for a better representation of the nuclear morphology data. Collectively, these data show that cytoskeletal disruption has only very small effects on nuclear shape.

