## Supplementary Materials for "Nuclear damage in *LMNA* mutant iPSC-derived cardiomyocytes is associated with impaired lamin localization to the nuclear envelope"

### Supplementary Information

#### Supplementary Figures

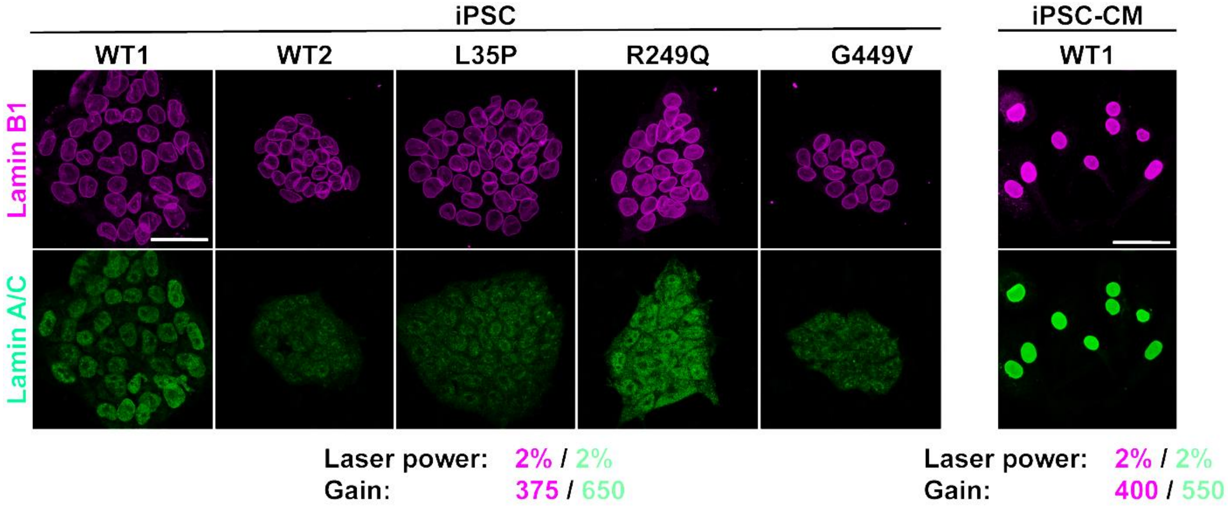


**Supplementary Figure 1.** iPSCs express low but detectable levels of Lamin B1 and Lamin A/C. Representative immunofluorescence images of WT and *LMNA*-mutant iPSCs, revealing detectable expression of Lamin A/C and Lamin B1. Expression of Lamin A/C was low compared to differentiated iPSC-CMs as expected, since Lamin A/C is upregulated with cardiac differentiation. Scale bars = 50 µm.

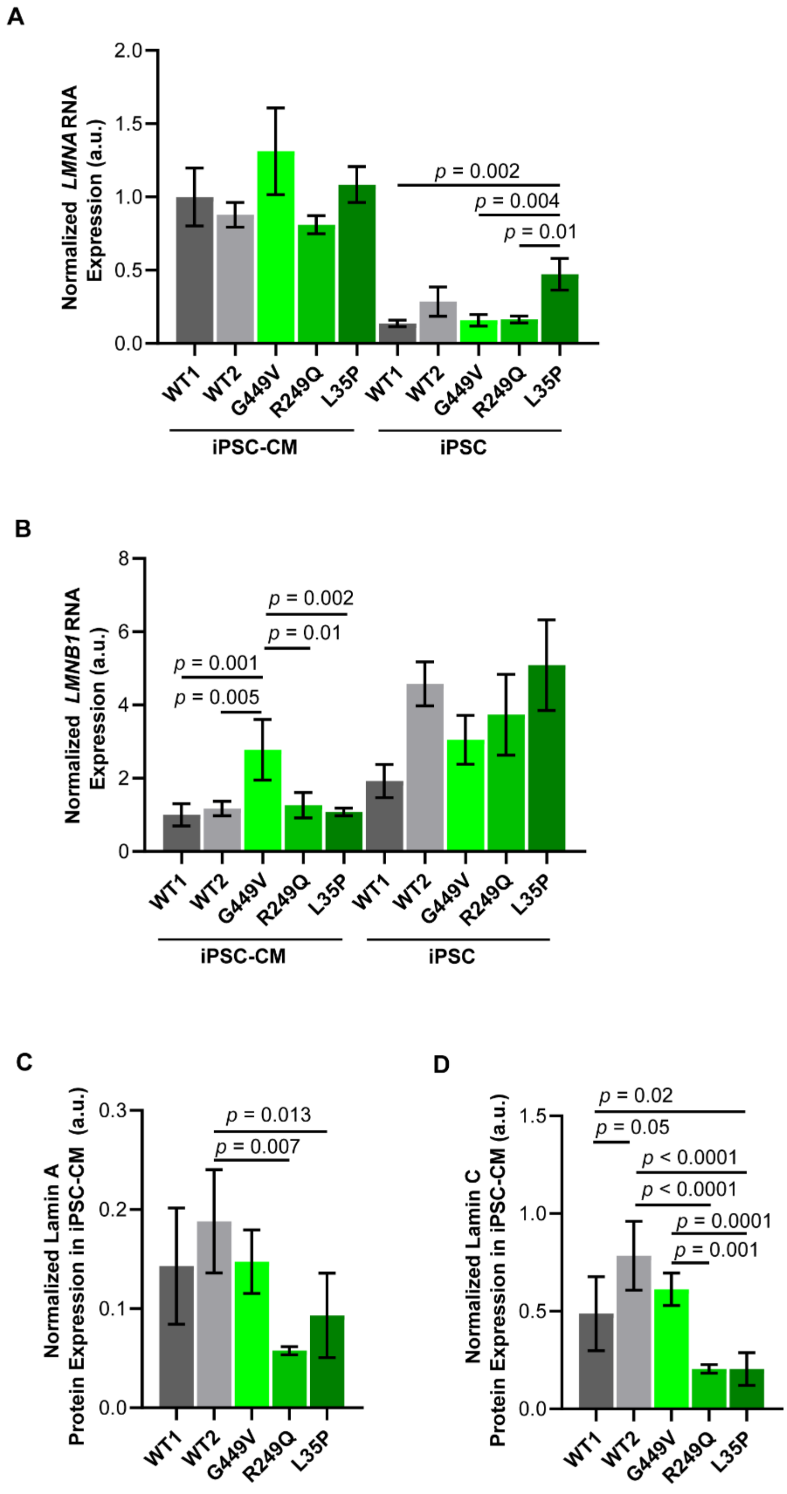


**Supplementary Figure 2.** *LMNA* mutant iPSCs and iPSC-CMs show normal levels of Lamin mRNA expression. (A) *LMNA* expression was low in iPSCs and increased with cardiac differentiation into iPSC-CMs. The difference in *LMNA* mRNA expression in the G449V iPSC-CMs was not statistically different compared to healthy controls iPSCs, and while L35P iPSCs exhibited increased *LMNA* expression compared to one healthy control, this difference did not carry over with cardiac differentiation. (B) *LMNB1* expression showed a general decrease with cardiac differentiation, and G449V iPSC-CM *LMNB1* expression was significantly increased compared to healthy controls. (C) Quantification of Lamin A protein expression from western analysis showed that L35P and R249Q iPSC-CMs had significantly decreased Lamin A protein expression compared to one healthy control line. (D) Quantification of Lamin C protein expression from western analysis showed that L35P and R249Q PSC-CMs had significantly decreased Lamin C protein expression compared to both healthy controls. Data presented as mean ± SEM. $N$ = 3-4 samples for all groups for mRNA levels. $N$ = 5 samples for all groups for western quantification.

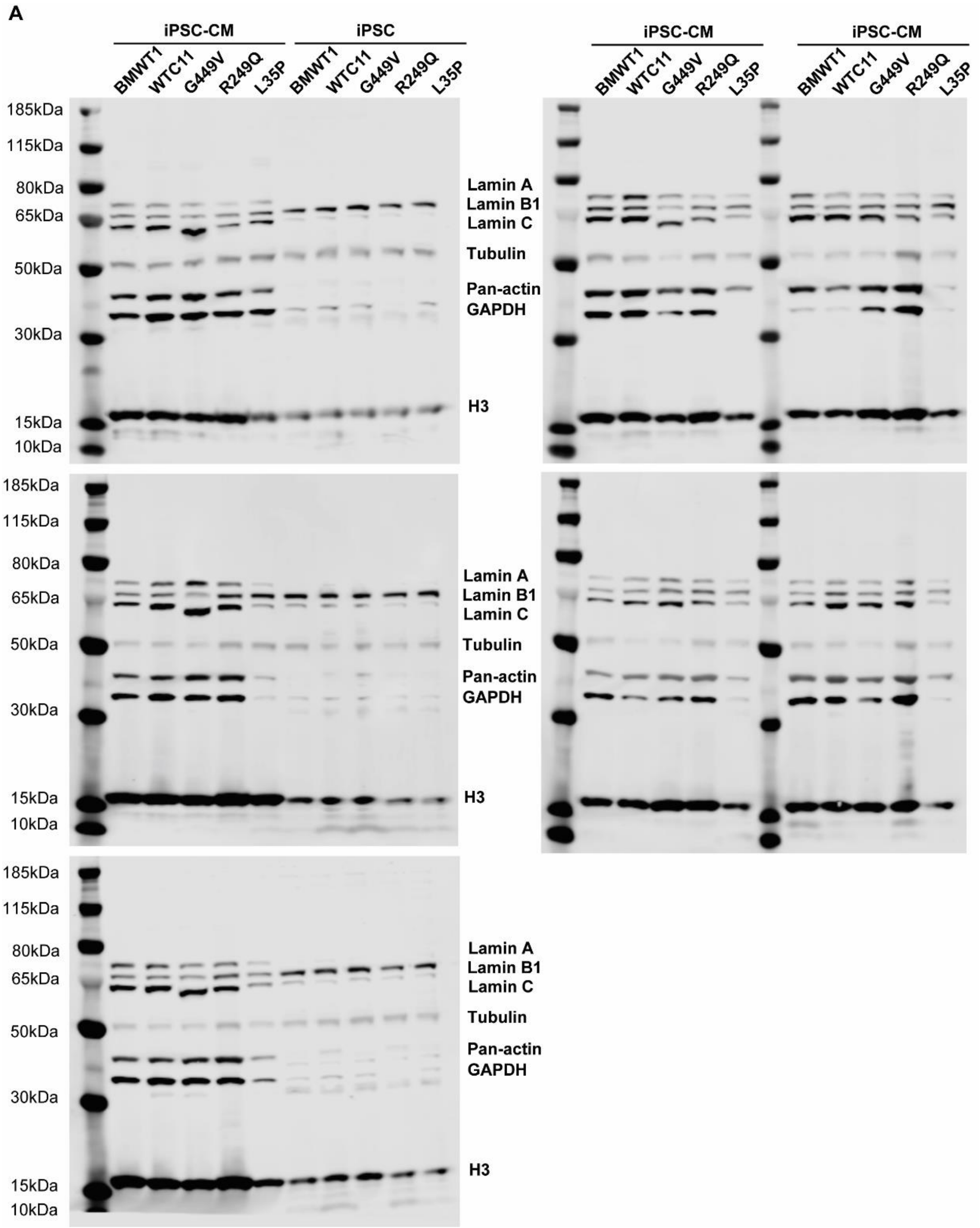


**Supplementary Figure 3.** Full immunoblots for quantification of lamin levels. (A) A total of five immunoblots each with one replicates per cell line and differentiation state (iPSC and iPSC-CM) were used to quantify protein levels of Lamin A/C and Lamin B1.

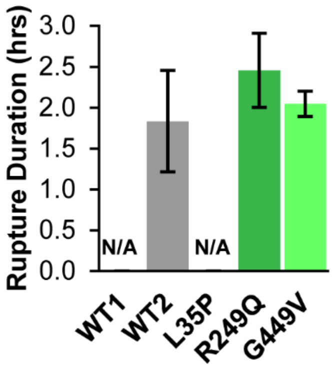


**Supplementary Figure 4.** Duration of nuclear envelope rupture is not altered by *LMNA* mutations. Quantification of nuclear envelope rupture duration, based on the time NLS-GFP was displaced from the nucleus, in *LMNA* mutant and healthy control iPSC-CMs. Difference in nuclear envelope rupture duration between healthy control (WT) and LMNA-mutant iPSC-CMs exhibiting nuclear envelope rupture were not statistically significant. Data presented as mean ± SEM. $N \geq 1020$ nuclei per group.

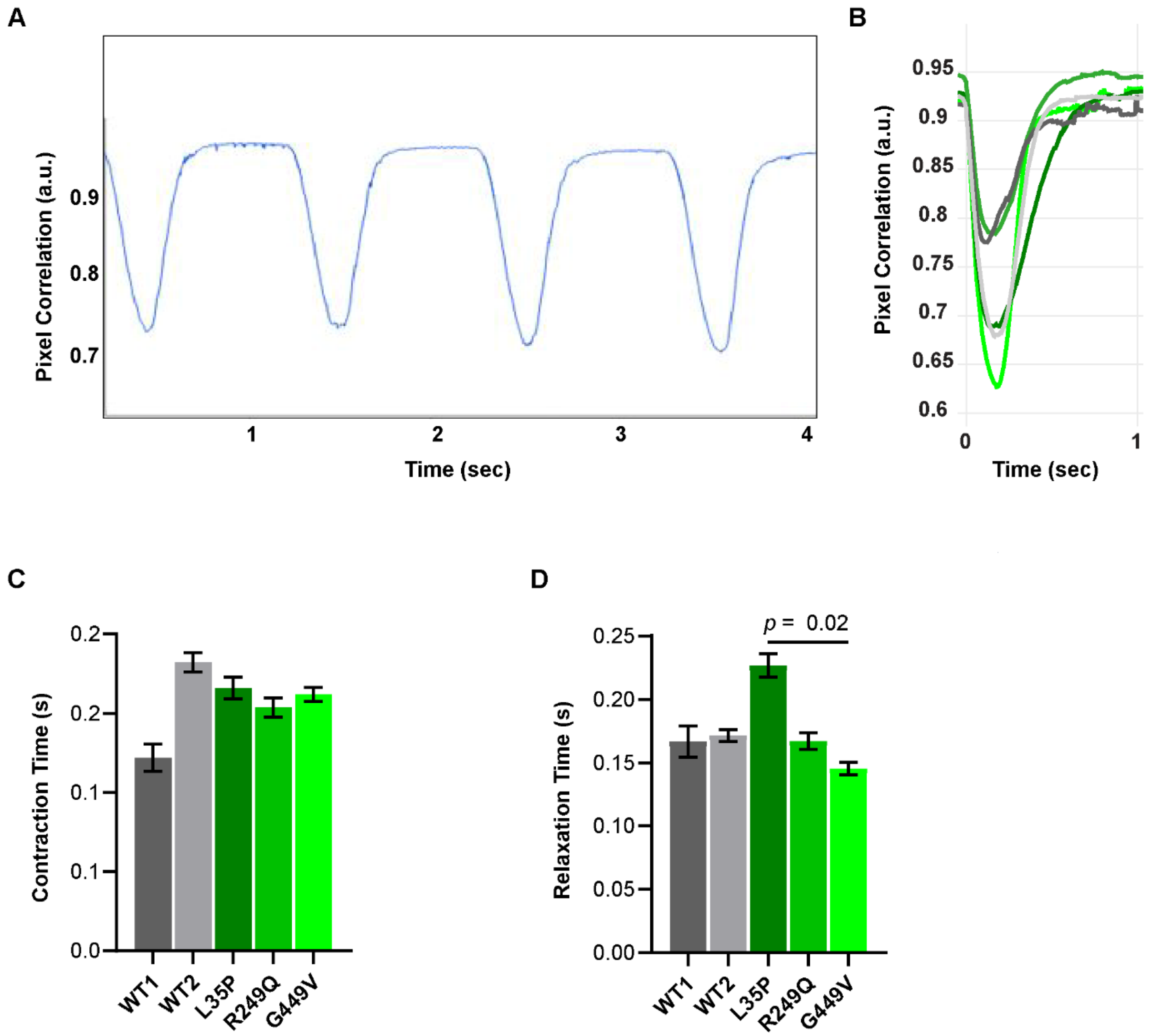


**Supplementary Figure 5.** *LMNA*-mutant iPSC-CMs do not exhibit altered contractility. (A) A representative contractile trace for WT2 iPSC-CM. (B) Averaged contraction trace per iPSC-CM cell line. (C) There are no significant differences in contraction time between healthy control and *LMNA* mutant iPSC-CM lines. (D) While L35P iPSC-CMs show significantly increased relaxation time compared to G449V iPSC-CMs, no *LMNA*-mutant iPSC-CM lines have significant changes in relaxation time compared to two healthy controls. Data presented as mean ± SEM. $N > 49$ cells per group from 3-4 independent repeats per cell line.

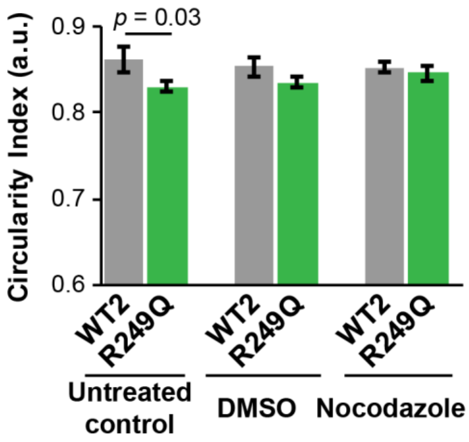


**Supplementary Figure 6.** Non-normalized nuclear circularity data for wild-type (WT2) and R249Q iPSC-CMs treated with either Nocodazole or vehicle control (DMSO), and untreated controls. Differences in the Nocodazole and DMSO treated groups are not statistically significant. Note that in contrast to the normalized data presented in Figure 5F, the effect of Nocodazole treatment on nuclear shape in the R249Q cells is not statistically significant, due to the overall increased variability in the non-normalized data. Data presented as mean ± SEM. $N \geq 44$ nuclei per group over two to three independent experiments for nuclear area and circularity index.

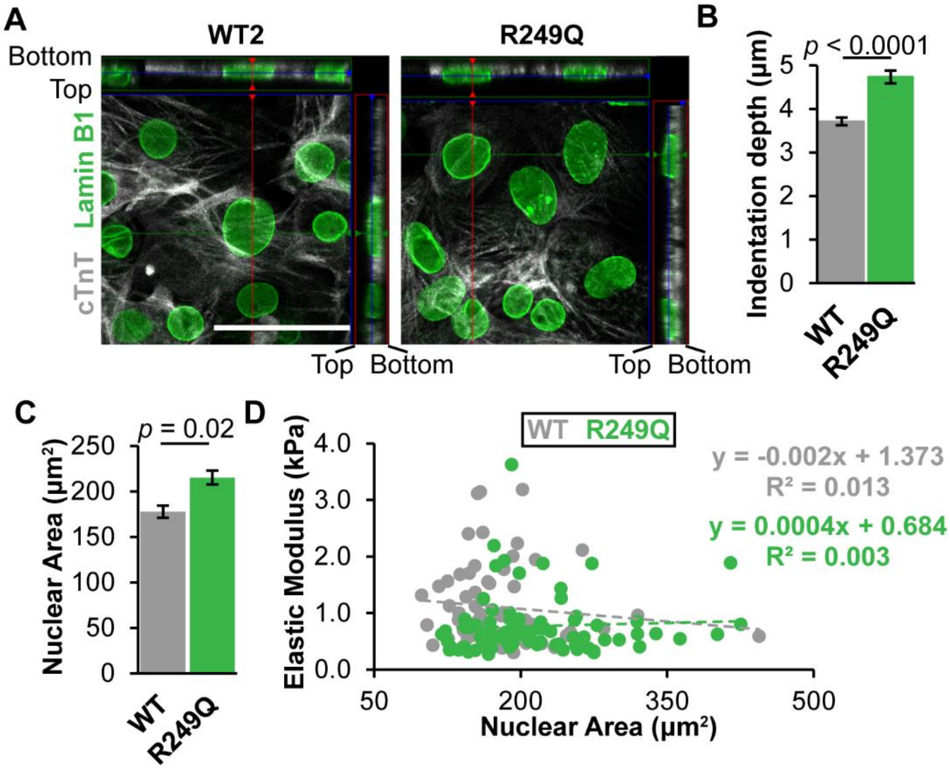


**Supplementary Figure 7.** R249Q iPSC-CM nuclear stiffness does not correlate to increased nuclear area. (A) An orthogonal view of representative healthy control (WT2) and R249Q iPSC-CMs shows that nuclei (labeled for Lamin B1, green) are almost directly exposed to the apical plasma membrane with very little cytoskeleton atop the nucleus, as indicated by immunofluorescence labeling for cTnT. Scale bar = 50μm. (B) Nuclear indentation depth was significantly increased in R249Q-iPSC-CM cells compared with WT2. These depths suggest that indentations were approximately 33-50% of the height of nuclei, as indicated in Figure 3D, and significantly more than the height of any cytoskeletal elements over the top of nuclei in panel A. (C) R249Q iPSC-CMs used in nuclear stiffness experiments had increased nuclear area, but (D) a scatter plot shows no correlation between the nuclear elastic modulus vs. nuclear cross-sectional area across individual cells. Equations indicate the results of linear regression models for each cell line. Data in B and C presented as mean ± SEM. $N \geq 68$ nuclei per group.

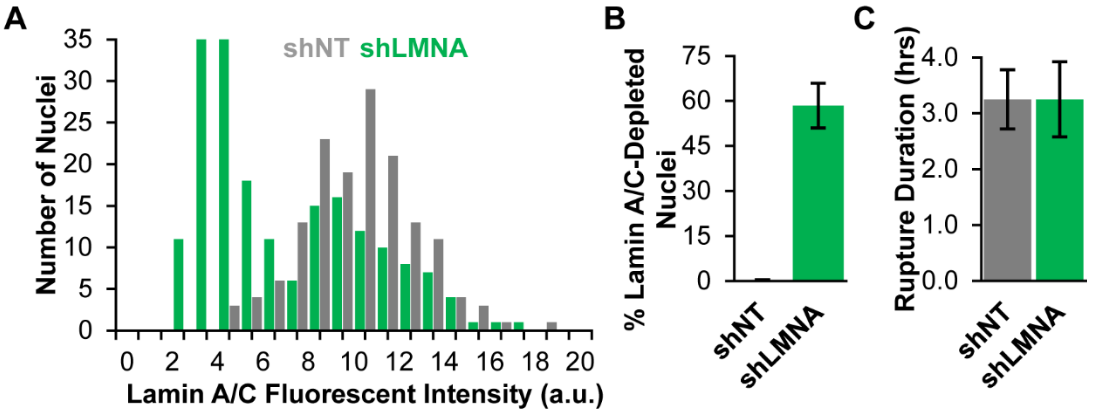


**Supplementary Figure 8.** Validation of shRNA mediated Lamin A/C depletion in healthy control iPSC-CMs. (A) Histogram of average nuclear Lamin A/C immunofluorescence intensities shows an increased population of nuclei with low or absent Lamin A/C expression in iPSC-CMs modified with shRNA targeting *LMNA* (shLMNA) or a non-target (shNT) control. Cells did not undergo selection prior to analysis. (B) Quantification of the number of nuclei with Lamin A/C Knockdown (KD) indicates >55% knockdown efficiency in the shLMNA treated cells. (C) shNT and shLMNA treated iPSC-CMs had similar durations of nuclear envelope rupture. Data in B and C presented as mean ± SEM. *N* > 150 nuclei per group for fluorescence intensity and Lamin A/C depletion efficiency, *N* > 1,415 nuclei per group for nuclear envelope rupture.

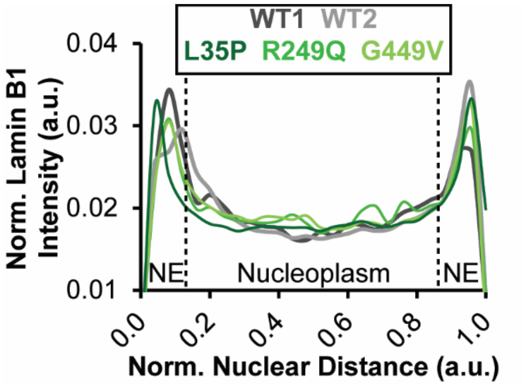


**Supplementary Figure 9.** Lamin B1 fluorescence intensity profiles. Lamin B1 fluorescence intensity profiles taken at the central z-plane of nuclei from confocal image sections of healthy control and *LMNA*-mutant iPSC-CMs.

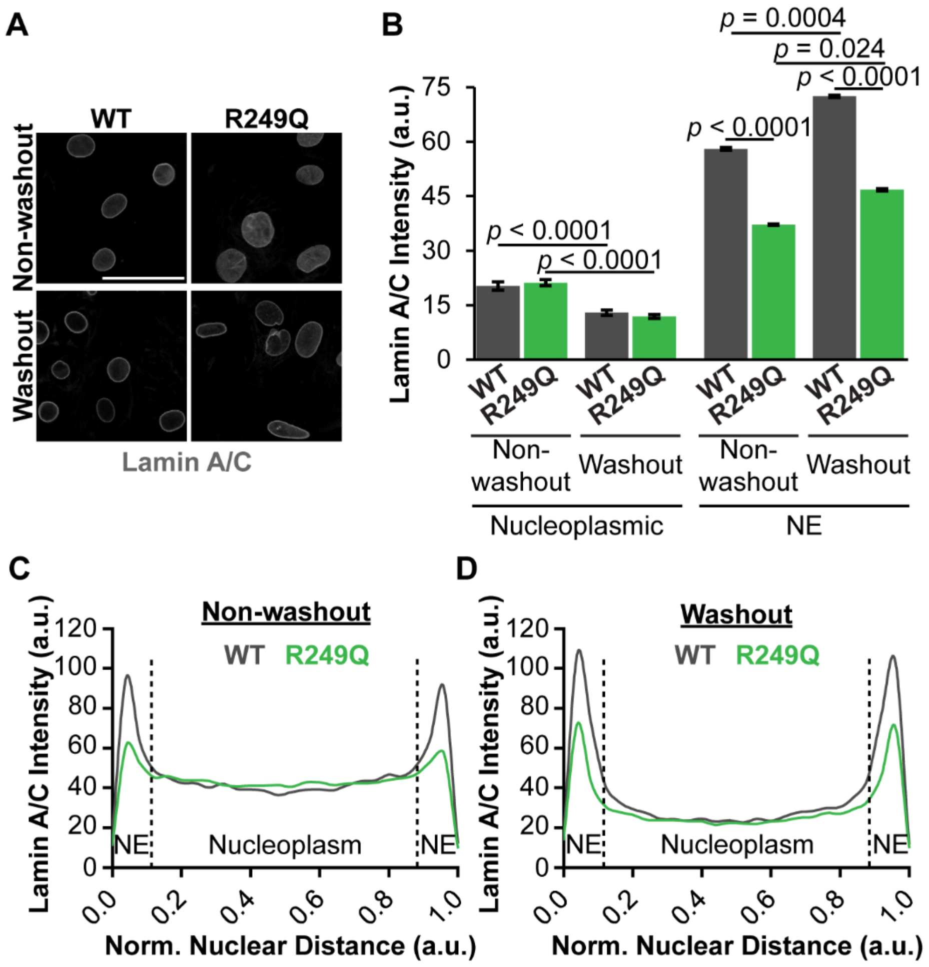


**Supplementary Figure 10.** *LMNA*-mutant iPSC-CMs have decreased polymerized Lamin A/C at the nuclear periphery, but normal levels of nucleoplasmic Lamin A/C. (A) Representative immunofluorescence images of Lamin A/C in healthy control (WT) and R249Q mutant iPSC-CMs following washout of soluble nuclear proteins and in non-washout controls. Scale bars = 50µm. (B) Quantification of the average nucleoplasmic and peripheral fluorescence intensities for healthy control (WT) and R249Q samples after washout and non-washout control showed that nucleoplasmic Lamin A/C levels decreased following washout but remained similar between cell lines. In contrast, R249Q iPSC-CMs had significantly reduced levels of peripheral Lamin A/C levels compared to healthy controls, and the differences remained after washout, indicating that the R249Q cells had reduced levels of assembled Lamin A/C at the nuclear lamina. These results are also visible in the Lamin A/C fluorescence intensity profiles of non-washout controls (C) and after washout (D). Data in B presented as mean ± SEM. $N > 60$ nuclei per group.

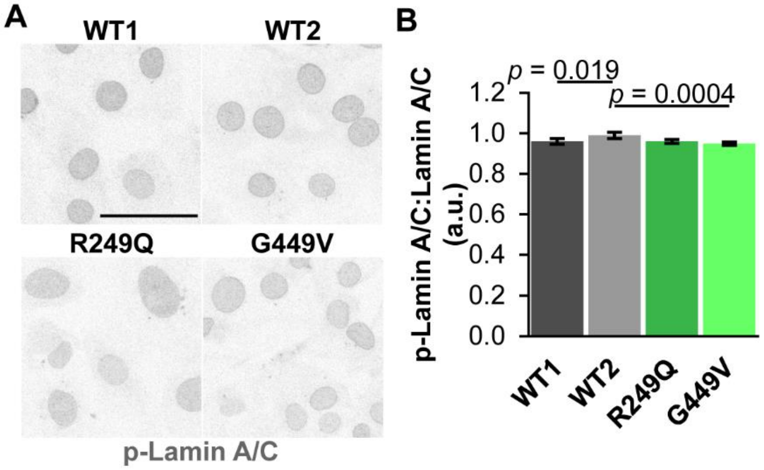


**Supplementary Figure 11.** *LMNA*-mutant iPSC-CMs do not have increased Lamin A/C phosphorylation. (A) Representative immunofluorescence images of *LMNA* mutant and healthy control iPSC-CMs immunofluorescently labeled for phospho-Ser22 Lamin A/C (p-Lamin A/C). Scale bar = 50µm. (B) The ratio of phospho-Lamin A/C to total Lamin A/C is not increased in *LMNA*-mutant iPSC-CMs. Data presented as mean ± SEM. $N > 60$ nuclei per group.

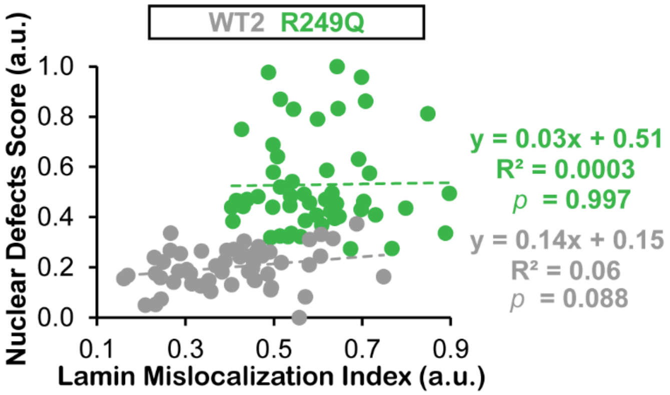


**Supplementary Figure 12.** Lamin mislocalization does not correlate to nuclear damage across single cells within individual iPSC-CM lines. Correlating lamin mislocalization index and nuclear defects score on individual nuclei for WT iPSC-CMs, which exhibit a lower degree a lamin mislocalization and nuclear damage, showed only a weak positive correlation ($R^2 = 0.06$), which was not quite statistically significant ($p = 0.088$). The R249Q nuclei did not show any correlation between lamin mislocalization and nuclear defects. $N > 50$ nuclei per cell line for single cell analysis.

**Supplementary Table(s)**

| Antibody | Catalog # | Vendor | Dilution (IF) | Dilution (Western) |
|---|---|---|---|---|
| Lamin A/C (E-1) | sc-376248 | Santa Cruz | 1:150 | 1:1000 |
| Lamin B1 | 12987-1-AP | Proteintech | 1:200 | 1:5000 |
| cTnT | RV-C2 | DSHB | 1:20 | |
| Phospho-Lamin A/C (Ser22) | 2026S | Cell Signaling | 1:500 | |
| Pan-Actin (D18C11) | 8456S | Cell Signaling | | 1:1000 |
| GAPDH | 60004-1-Ig | Proteintech | | 1:20,000 |
| H3 | | Abcam | | 1:5000 |
| Alexa Fluor 488; Goat anti-mouse IgG1 | A-21121 | Invitrogen | 1:250 | |
| Alexa Fluor 568; Goat anti-mouse IgG1 | A-21124 | Invitrogen | 1:250 | |
| Alexa Fluor 647; Goat anti-mouse IgG2b | A-21242 | Invitrogen | 1:250 | |
| Alexa Fluor 488; Donkey anti-rabbit IgG | A-21206 | Invitrogen | 1:250 | |
| Alexa Fluor 568; Donkey anti-rabbit IgG | A-10042 | Invitrogen | 1:250 | |
| IRDye 800CW Donkey anti-Rabbit IgG | 926-32213 | Li-cor | | 1:3000 |
| IRDye 880RD Donkey anti-Mouse IgG | 926-68072 | Li-cor | | 1:3000 |

**Supplementary Table 1.** Primary and secondary antibodies used for immunofluorescence labeling and western analysis.

| Primer | Forward (5'-3') | Reverse (5'-3') |
|---|---|---|
| *h-LMNA* | GACTCAGTAGCCAAGGAGCG | TTGGTATTGCGCGCTTTCAG |
| *h-LMNB1* | AAGCAGCTGGAGTGGTTGTT | TTGGATGCTCTTGGGGTTC |
| *h-GAPDH* | TGCGTCGCCAGCCGAG | AGTTAAAAGCAGCCCTGGTGA |
| *h-TMX4* | GGGTCTGGTCTTGGTGGTAA | AGCCTCCTCTGATCTCCGATT |
| *h-18S* | CTCAACACGGGAAACCTCAC | CGCTCCACCAACTAAGAACG |

**Supplementary Table 2.** Primers used for qPCR analysis.
